# Genetic-variant hotspots and hotspot clusters in the human genome facilitating adaptation while increasing instability

**DOI:** 10.1101/2020.10.16.342188

**Authors:** Xi Long, Hong Xue

**Affiliations:** Division of Life Science and Applied Genomics Centre, Hong Kong University of Science and Technology, Clear Water Bay, Hong Kong, China; HKUST Shenzhen Research Institute, 9 Yuexing First Road, Nanshan, Shenzhen, China; Centre for Cancer Genomics, School of Basic Medicine and Clinical Pharmacy, China Pharmaceutical University, Nanjing, Jiangsu, China

**Author notes:** Author for Correspondence: Hong Xue, Division of Life Science and Applied Genomics Centre, Hong Kong University of Science and Technology, Clear Water Bay, Hong Kong, China.

**Keywords:** Frequency independent, Genetic diversity, Missing heritability, Retrotransposon, Recombination-selection co-saturation, Replication timing

## Abstract

**Background:** Genetic variants, underlining phenotypic diversity, are known to distribute unevenly in the human genome. A comprehensive understanding of the distributions of different genetic variants is important for insights into genetic functions and disorders.

**Methods:** Herein, a sliding-window scan of regional densities of eight kinds of germline genetic variants, including single-nucleotide-polymorphisms (SNPs) and four size-classes of copy-number-variations (CNVs) in the human genome has been performed.

**Results:** The study has identified 44,379 hotspots with high genetic-variant densities, and 1,135 hotspot clusters comprising more than one type of hotspots, accounting for 3.1% and 0.2% of the genome respectively. The hotspots and clusters are found to co-localize with different functional genomic features, as exemplified by the associations of hotspots of middle-size CNVs with histone-modification sites, work with balancing and positive selections to meet the need for diversity in immune proteins, and facilitate the development of sensory-perception and neuroactive ligand-receptor interaction pathways in the function-sparse late-replicating genomic sequences. Genetic variants of different lengths co-localize with retrotransposons of different ages on a ‘long-with-young’ and ‘short-with-all’ basis. Hotspots and clusters are highly associated with tumour suppressor genes and oncogenes (*p* < 10^−10^), and enriched with somatic tumour CNVs and the trait- and disease-associated SNPs identified by genome-wise association studies, exceeding tenfold enrichment in clusters comprising SNPs and extra-long CNVs.

**Conclusions:** In conclusion, the genetic-variant hotspots and clusters represent two-edged swords that spearhead both positive and negative genomic changes. Their strong associations with complex traits and diseases also open up a potential ‘Common Disease-Hotspot Variant’ approach to the missing heritability problem.

## Background

Genetic variants (GVs) are essential contributors to population diversity, providing an important basis for the investigation of genetic effects on complex traits and disease susceptibility. A wide spectrum of GVs exist in the human genome ranging from point alterations, viz. single-nucleotide-polymorphisms (SNPs), to structural variations including copy-number-variations (CNV). It was found that a typical human genome differed from the reference genome at a median of 4.31 million base pairs, 84.5% of which consisted of SNPs, the most common form of genetic variants in human genomes [1]. Although occurring at lower frequencies, structural variations contribute significantly to genetic diversity among individuals and populations by causing sequence alternations ranging from several to millions of base pairs.

GVs have been analyzed in genome-wide association studies (GWAS), revealing hundreds of thousands of associations of GVs with complex diseases and traits [2]. SNPs are often employed as genetic markers in the ‘Common Disease-Common Variant’ hypothesis-based GWAS [3–5] with the rationale that common diseases may be expectedly attributable to common genetic variants [6–9]. However, so far only a small fraction of SNP-disease or SNP-trait associations have been discovered, representing a potential component of the missing heritability regarding complex disorders and traits [7, 10, 11]. Other contributing factors to missing heritability would include the poor detection of rare variants strongly associated with complex diseases or traits, giving rise to the ‘Common Disease-Rare Variant’ hypothesis [8, 12] which has been confirmed by discoveries of disease- or trait-associated rare structural variants such as CNVs [13–15]. Additionally, a ‘hypothesis driven’ approach is found to be useful in examples such as the identification of Autism-associated SNPs where shift of codon usage is postulated to alter protein translation efficiency [16, 17]. Models also have been developed to delineate the effects of demographic history and genetic forces on the patterns and maintenance of genetic variants underlining complex traits [18–20]. Nonetheless, a substantial portion of the heritability remains unexplained, giving rise to the need for a more comprehensive understanding of the distribution, formation mechanisms and functional effects of different kinds of genetic variants as a possible gateway to a further reduction of this unexplained portion.

Previous studies have indicated that genetic variants are not evenly distributed in the human genome [21, 22], with density enhancements giving rise to genetic-variant hotspots in some genomic regions, and identified homologous recombination as one of the major mechanisms for the formation of GV hotspots [23–25]. Since transposable elements constitute highly abundant repeat sequences in the genome, they can serve as homologous templates in recombination events that produce GVs [26–28]. Rapid accumulation of sequence variations has been observed in the body of young Alu elements [29], erasing the sequence similarity between homologous Alu pairs, and reducing thereby their capability to serve as templates for homologous recombination. It follows that the age of transposable elements could be associated with different types and frequencies of GVs. Genetic variants could also originate from mechanisms such as background mutations, non-homology repairs and replicative errors [30–32]. In addition, the choice of DNA double-strand break repair pathways has been found to be cell cycle-dependant [33, 34], making replication timing an important factor in shaping the distribution of GVs.

Human CNV hotspots have been found to overlap with CNVs in the chimpanzee and macaque genomes, pointing to the maintenance of these hotspots by nonneutral evolutionary forces [24]. Such forces could underlie the differential distribution of different kinds of genomic features and genetic variants among the three types of sequence zones in the human genome, viz. the Genic (gene-rich), Proximal (gene-proximal) and Distal (gene-distal) zones [35]. Accordingly, the present study is directed to establishing a high-resolution comprehensive landscape of genetic-variant hotspots and hotspot clusters in the three types of sequence zones, so that an improved understanding may be obtained regarding their formation mechanisms, and constraint forces, their adaptive or destabilizing roles in the genome, and their possible relevance to the problem of missing heritability.

## Materials and Methods

### Data sources

Germline genetic variants employed in genetic variant hotspots detection include biallelic SNPs (~ 77 million entries) and small indels (SIDs, ~ 3 million) from the Phase III 1000 Genomes Project [1], CNVs (~ 1.6 million) from dbVar database [36], microsatellites (MSTs, ~ 40,000) from the ‘Microsatellite track’ [37], and segmental duplication (SDPs, ~ 50,000) from the ‘Segmental Dups track’ [38] of the UCSC Genome Browser database [39]. Rare SNPs (minor allele frequency < 1%) located within the exome pull-down target boundaries are excluded from analysis on account of the probability of the rare alleles discovered given disproportionally high coverage of the exons and their flanking regions in the 1000 Genomes Project. The data sources of the 55 functional and structural genomic features analyzed in the present study are given in the ‘Data source and track detail’ column of Supplementary Table S1A. Information of the germline genetic variants are available in Supplementary Table S1B. The genomic coordinates of gap regions of human genome assembly hg19 are retrieved from the ‘Gap’ track of the UCSC Genome Browser database. Genomic coordinates of immune system gene loci are retrieved from NCBI Gene database [40].

### Size-classifications of CNVs

The length distribution of germline CNVs are separated into four size classes using cut points determined as the critical points of the polynomial regression curve fitted to two successive peaks on the length distribution curve as shown in Supplementary Figure S1. The four size classes comprise the short CNVs (5 < length ≤ 61 base pairs [bp]), medium CNVs (61 < length ≤ 952 bp), long CNVs (952 < length ≤ 15,571 bp) and extra-long CNVs (length > 15,571 bp).

### Genomic feature quantitation and tripartite genomic zones

For a feature where the number of base pairs is counted, its level in any region is expressed in ‘Density’, e.g. for the level of CpG islands in a certain region:

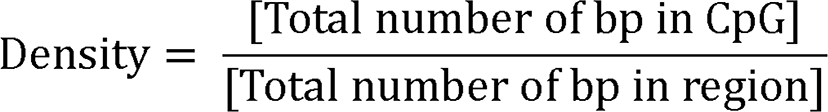

On the other hand, for a feature that is assessed by a numerical score, its level is expressed in ‘Intensity’, e.g. for the level of methylation in any genomic region:

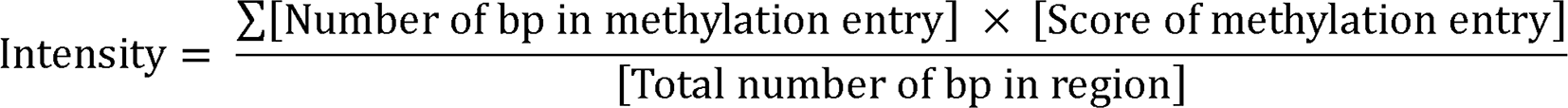

Various features expressed in ‘Density’ or ‘Intensity’ are listed in ‘Density/Intensity’ column of Supplementary Table S1. Genomic features are classified into the Genic, Proximal and Distal groups based on their co-localization patterns in 500-kb successive and non-overlapping sequence windows in the twenty-two autosomes (Supplementary Figure S2); based on their feature compositions, the 500-kb windows are partitioned into 45.1% Genic, 31.1% Proximal and 23.8% Distal zone windows as described by Ng et al. [35] covering 94.0% of total non-gap sequences on the 22 autosomes. Accordingly, autosomal regions in the present study refer to genomic sequences that have been assigned to the Genic, Proximal, or Distal zone windows on the 22 autosomes.

### Density-based genetic-variant hotspots determined by weighted sliding windows

To increase the resolution of hotspot detection, a sliding-window protocol is employed to scan the genome in 1-kb windows sliding by 10-bp steps. As well, to render more precise the genomic boundary of any hotspot as illustrated in Supplementary Figure S3, the number of genetic variant entries in each of the one hundred 10-bp steps (or, n_step_) in a window is assigned a weight w_step_ that depends on its distance from the window centre, increasing in equal increments from a w_step_ of 0.5 at the extreme margins of the window to 1.0 at the very centre of the window. This gives rise to a symmetrical distribution of step weights within the window, and the weighted density of any genetic variant in each window (D_win_) is given by:

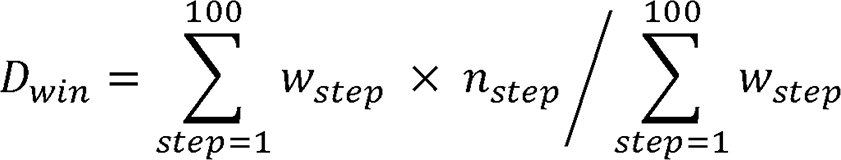

The weighted density of the targeted kind of genetic variants is calculated in all 1-kb windows. In the Genic, Proximal or Distal zones, the top-density windows that cover altogether 5% of the total entries of the targeted genetic variant are identified as hotspots. For example, the hotspot detection procedure for SNPs in the Genic zones (see flowchart in Supplementary Figure S4) consists of: (i) each autosome of the human genome is divided into successive 1-kb sliding-windows in 10-bp steps; (ii) D_win_ for SNP entries is determined for all the sliding windows; (iii) those sliding windows that overlap with a 500-kb Genic zone are separated into Part I sliding windows that are entirely located within a Genic zone, and Part II sliding windows that are partially located within a Genic zone; (iv) Part I and Part II windows are ranked separately based on their D_win_ values from the highest to the lowest. The Part I sliding windows with top-ranked D_win_ values are recruited successively from rank 1 down to rank i until their cumulative SNP fraction represents ≤ 5% of total SNP entries in the Genic zones to yield the top SNP windows in Part I (with ranks of 1, 2, 3, … i, red-numbered in the flowchart). The top-ranked Part II sliding windows are recruited from rank 1 down to rank k (with ranks of 1, 2, 3, … k, red-numbered in flowchart) such that the rank-k window displays a weighted SNP density (D_k_) greater than or equal to the D_min_ (viz. lowest D_win_) among the top-ranked SNP windows in Part I. Steps iii and iv are also repeated for the Proximal and Distal zones as described for the Genic zones. (v) Thereupon, Part I and Part II top-ranked windows from the three types of zones are merged together, followed by elimination of any double-counting of windows on account of their zone-crossing to yield altogether a total of 34,487 SNP hotspots in the three types of zones with a mean width of 1,994 ± 4,528 bp, covering 2.54% of the autosomal regions analyzed. The zone-specific hotspot detection thresholds (D_min_), and the chromosomal coordinates of different kinds of genetic variant hotspots, are given in Supplementary Table S2 and Supplementary Dataset S1.

### Identification of hotspot clusters

When two or more of different kinds of genetic-variant hotspots overlap with one another, the genomic regions occupied by these hotspots are merged into a hotspot cluster. This procedure identifies 1,135 clusters in the genome consisting of 2 to 3 kinds of hotspots each, amounting to 0.20% of autosomal sequences and comprising twenty-three kinds of hotspot-compositions, the chromosomal coordinates of which are given in Supplementary Dataset S1.

### Determination of natural selection hotspots

To identify positive selection hotspots (PosSel-Hs), the strength of positive selection is evaluated in 1-kb windows in the non-gap regions on 22 autosomes based on i) average derived-allele frequency (DAF) of all SNPs in each window, ii) average of the maximum DAF differences (|ΔDAF|) across the two different populations, and iii) the average of haplotype structure-based statistic |nS_L_| [41]. Next, the top-5% windows ranked according to each of these three kinds of positive-selection levels are earmarked as candidate windows under positive selection. All 1-kb windows in the autosomes are classified into ten ranking groups based on their number of informative sites (viz. the SNPs with DAF or nSL statistics). For each candidate window, its level of positive selection is compared with the estimated levels in 10,000 random windows simulated from the corresponding ranking group. The matching of informative site density would accommodate the higher variance in site-based measurement in the windows with low numbers of informative sites. Only windows with significantly higher measurement of positive selection (*p* < 0.05) relative to the random simulations are kept, and successive windows are merged to yield the positive selection hotspots. In this regard, 99,266 DAF hotspots, 89,464 |ΔDAF| hotspots, and 63,060 |nS_L_| hotspots are identified, which altogether cover 10.33% of autosomal regions. Likewise, the negative selection hotspots (NegSel-Hs) are identified based on two different measures of purifying selection, viz. i) sequence conservation score measured using phyloP across 100 species, and ii) intensity of nucleotide diversity. The windows with top-5% phyloP score or bottom-5% nucleotide diversity are earmarked, and only the windows displaying significantly higher measurements of purifying selection (*p* < 0.05) relative to 10,000 simulation windows of similar levels of informative sites are regarded as NegSel-Hs, which altogether amount to 7.94% of autosomal sequences. The phyloP score is retrieved from the ‘Conservation’ track of UCSC Table Browser [42]. The nucleotide diversity in any window is estimated using VCFtools version 0.1.15 with the ‘--window-pi’ flag [43]. The |nS_L_| value for each SNP is the absolute value of the nS_L_ statistic estimated and normalized across 100 frequency bins in selscan version 1.2.0 with all the default settings [41, 44]. The DAF, |ΔDAF|, |nS_L_| and nucleotide diversity used for selection-hotspot determination are based on SNPs and haplotypes from the Phase III 1000 Genomes Project of 2,504 individuals. Genomic windows subject to balancing selection retrieved from the Supplementary Material of Bitarello et al. [45] are merged across African and European populations at different target frequency values, yielding 10,275 balancing selection hotspots (BalSel-Hs) amounting to 1.71% bp of the autosomal regions. Summaries of selection hotspots are available in Supplementary Table S3A. The chromosomal coordinates and *p*-values of each of the PosSel-Hs and the NegSel-Hs identified are given in Supplementary Dataset S1.

### Replication-time segments

Based on the previously assessed DNA replication times of 1-kb sequence windows in fifteen cell lines retrieved from the ‘UW Repli-seq track’ in the UCSC Table Browser [46], the human genome is classified into six types of replication-time segments in the present study as follows: (i) for each cell line, record the replication phase(s) that earns the highest score among the six replication phases (G1b, S1, S2, S3, S4 and G2 phases) within every 1-kb window; (ii) for any window, the replication phase that earns the highest score most frequently from the 15 cell lines is chosen as the representative replication phase for all the 15 cell lines. Only windows whose replication time has been assessed in more than eight out of the fifteen cell lines are subject to analysis, yielding 325,635 G1b, 389,304 S1, 395,223 S2, 454,554 S3, 392,600 S4 and 512,129 G2 1-kb windows that fall into the Genic, Proximal or Distal zonal windows covering altogether 85.7% of the twenty-two autosomes. Windows yielding more than one representative replication phases are not included in the analysis. Successive windows that share the same representative replication phase are merged to yield segments of varied lengths, and the chromosomal coordinates of the six types of replication-time segments are given in Supplementary Dataset S1.

### Somatic CNV breakpoints in cluster-containing genes

Coordinates of 23,056 known genes are matched with the locations of hotspot clusters to yield 448 genes that overlap with one or more clusters, and 442 of them display one or more CNVT breakpoint(s). For each of these 442 genes, the cluster-segments and noncluster-segments are compared with respect to their CNVT breakpoint densities, identifying 33 genes where the cluster-segments are significantly enriched in CNVT breakpoints (Chi-square test, Bonferroni-corrected), and only one gene where the cluster-segment is significantly depleted in CNVT breakpoints as shown in Supplementary Table S4. In the Chi-square tests, the expected number of CNVTs in the cluster-segment of the gene is calculated by multiplying the number of total CNVTs in the gene and the fraction of base pairs covered by the cluster-segments in the gene. Gene coordinates were retrieved from the R package ‘TxDb.Hsapiens.UCSC.hg19.knownGene’ version 3.2.2 using ‘genes’ function in ‘GenomicFeatures’ package version 1.26.4 [47].

### Statistical analysis

#### Empirical p-values based on Monte Carlo simulations

In each round of simulation, the locations of target regions are randomly shuffled (by matching the number and sizes of the target regions) within the autosomal regions analyzed. The measurement of interest is assessed for each simulation round. The *p*-value is obtained as (r+1)/(n+1) [48], where n is the round of simulations and r is the number of simulations that produce a measurement greater than or equal to the actual level in the case of significant enrichment, or less than or equal to the actual level in the case of significant depletion. Unless otherwise specified, 5,000,000 rounds of simulations are performed.

#### Enrichment of GWAS-identified SNPs in hotspots and clusters

To estimate using the Monte Carlo method the level of significance regarding the enrichment of GWAS-identified SNPs in the genetic-variant hotspots and hotspot clusters, random regions are simulated in sequence windows with matching levels of average minor-allele frequencies (MAF) in order to accommodate the dependency of GWAS on allele frequency. For this purpose, autosomal sequence windows are classified into ten equal groups according to their individual average MAF values, and the hotspots or clusters are also classified into the same ten groups. The Monte Carlo simulations are conducted by simulating sequence windows from the corresponding MAF group of hotspot or cluster. The simulation results for each group of hotspots and clusters are shown in Supplementary Figure S5.

#### Comparison of population differentiation between hotspot clusters and non-hotspot regions

To compare the positive-selection strength estimated by population differentiation among the cluster and non-hotspot (viz. the genomic regions outside of hotspots and clusters) regions, the DAF difference between any two populations, viz. |ΔDAF|^*POP1*-*POP2*^, is estimated for all the clusters and non-hotspot regions in each of the ten population pairs among African, American, South Asian, European and East Asian. Let 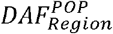 represents the average DAF of all SNPs in a specific cluster or non-hotspot region in a population. The |ΔDAF| between any two populations, for example African and European, viz. |ΔDAF|^*African*-*European*^, pertaining to the i^th^ cluster and j^th^ non-hotspot region is given by 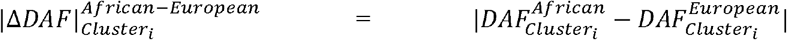 and 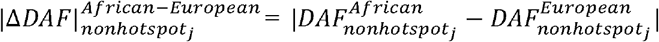. Thereupon, in the case of ***n*** total clusters and ***m*** total non-hotspot regions, the 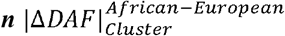 values are compared with the 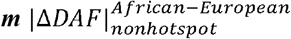 values by unpaired one-tailed t-tests. The procedure is likewise repeated for the other nine population pairs.

### Software employed in data processing and visualization

Custom R scripts are employed in hotspot and cluster detections and analyses under R environment version 3.3. Disease- or trait-associated SNPs identified by GWAS at *p* < 10^−8^ are lifted over from GRCh38.p5 to hg19 using the ‘liftOver” function in R package ‘rtracklayer’ version 1.34.2 [49]. Processing and quantification of genetic variants and genomic features are conducted using R package ‘GenomicRanges’ version 1.26.4 [47]. Figures are drawn using Cytoscape version 3.6.1 [50, 51], as well as the R packages ‘ggplot2’ version 3.0.0 [52], and ‘quantsmooth’ version 1.40.0.

## Results

### Density-based genetic variant hotspots and clusters

The germline genetic variants retrieved from different public databases are analyzed including SNPs, SIDs, MSTs, CNVs, as well as SDPs which originated from fixed CNVs [53] (see “Data source” in Materials and Methods). The three cuts on the length distribution of germline CNVs in Figure 1A separate the CNVs into four size classes comprising the short CNVs (SCNVs), medium CNVs (MCNVs), long CNVs (LCNVs) and extra-long CNVs (ECNVs). Notably, the presence of all four different size-classes of CNVs is evident in all three types of genomic zones (Figure 1A, panels 2-4).

**Fig. 1.**
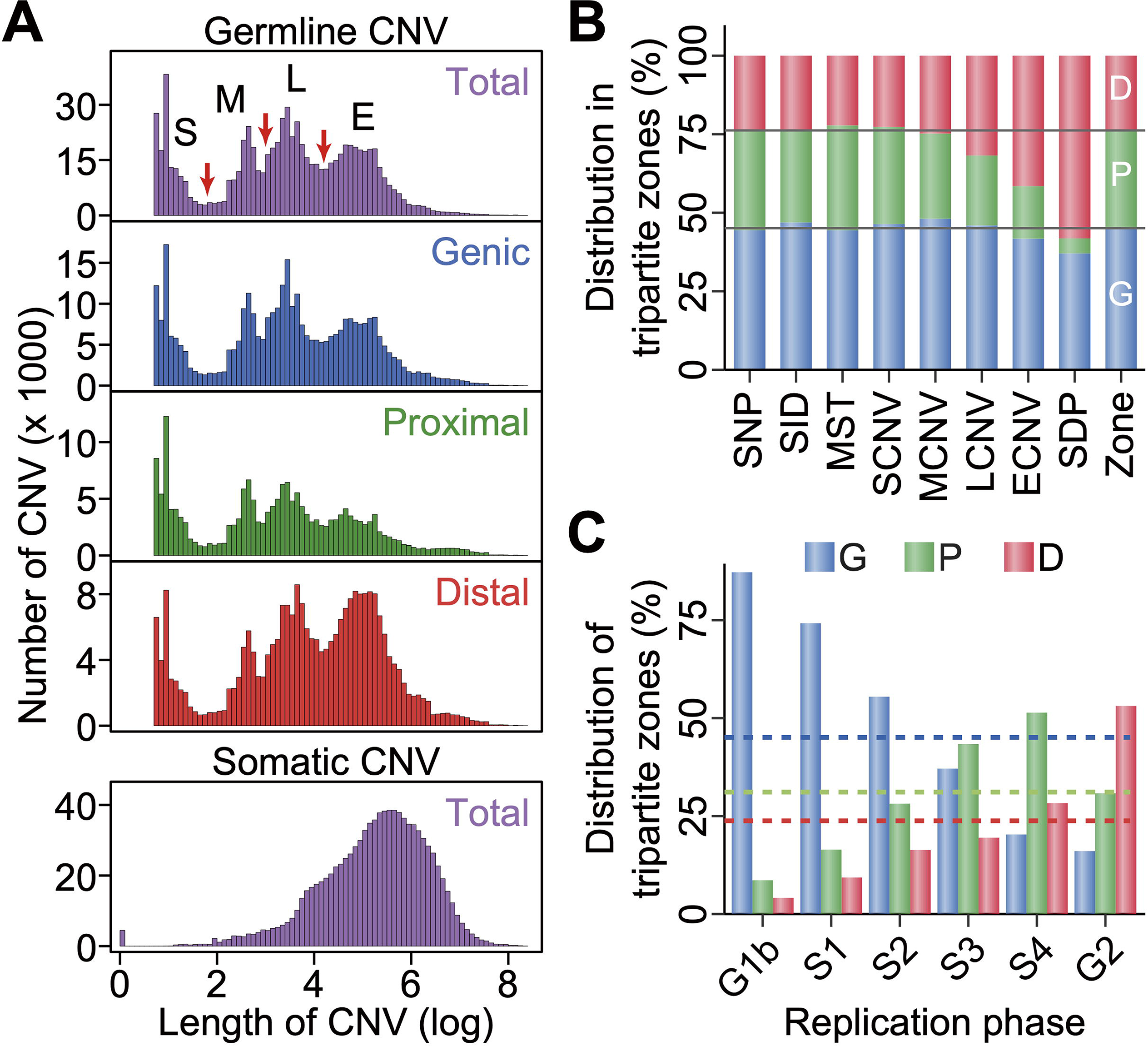
Genetic variants and replication phases in different genomic zones. **(A)** Length distributions of CNVs. Upper panel: distribution of germline CNVs in total and in the Genic, Proximal and Distal genomic zones. Germline CNVs are separated into short (S), medium (M), long (L) and extra-long (E) CNVs based on the three cuts indicated by the red arrows (see ‘Size-classifications of CNVs’ in Methods). Lower panel: distribution of total somatic CNVs in tumors from COSMIC database. **(B)** Percentile distributions of eight kinds of genetic variants in the Genic (blue), Proximal (green) and Distal (red) zones. The last ‘Zone’ column expresses the percentages of base pairs in the Genic, Proximal and Distal zones amounting to 45.1%, 31.1% and 23.8% respectively of the total base pairs of the human genome. **(C)** Percentages of Genic-, Proximal- and Distal-zone sequences in six different phases of DNA replication ranging from the earliest-replicating G1b to the latest-replicating G2 phases. Colored dashed lines indicate the 45.1% Genic-, 31.1% Proximal- and 23.8% Distal-zone sequences of the human genome. G, Genic; P, Proximal; D, Distal.

Because the average frequencies of genetic variants vary between the Genic, Proximal and Distal zones (Figure 1B), the identification of genetic-variant hotspots is performed separately for the three types of zones. To enhance resolution and obtain more precise boundary of genetic-variant hotspots, a weighted sliding-window algorithm with windows of 1,000 bp in steps of 10 bp is employed. For each type of genomic zones, the weighted density of any targeted genetic variant is measured in the sliding windows, and all the windows are ranked based on their densities from high to low. The top-density windows that cover altogether up to 5% of the total entries of the targeted genetic variant are identified as hotspots of that kind of genetic variant (see “Density-based genetic-variant hotspots determined by weighted sliding windows” in Materials and Methods). Altogether 44,379 hotspots with an average size of 1,882 bp are identified in the three genomic zones, equivalent to 3.09% of the total base pairs in the twenty-two autosomes (Figure 2A and Supplementary Table S2). The enrichments of different kinds of GVs in their respective hotspots relative to simulated genomic regions are significant with *p*-values ranging from 4 × 10^−4^ to 0 (Supplementary Figure S6). In general, the constitutive GVs in the hotspots display higher allele frequencies than genomic average except in the case of the SID hotspots (Supplementary Figure S7). On the basis that the convergence of two or more kinds of hotspots produces a hotspot cluster (see “Identification of hotspot clusters” in Materials and Methods), there are 1,135 clusters with an average size of 4,816 bp, equivalent to 0.20% of the total base pairs (Figure 2A and Supplementary Table S2).

**Fig. 2.**
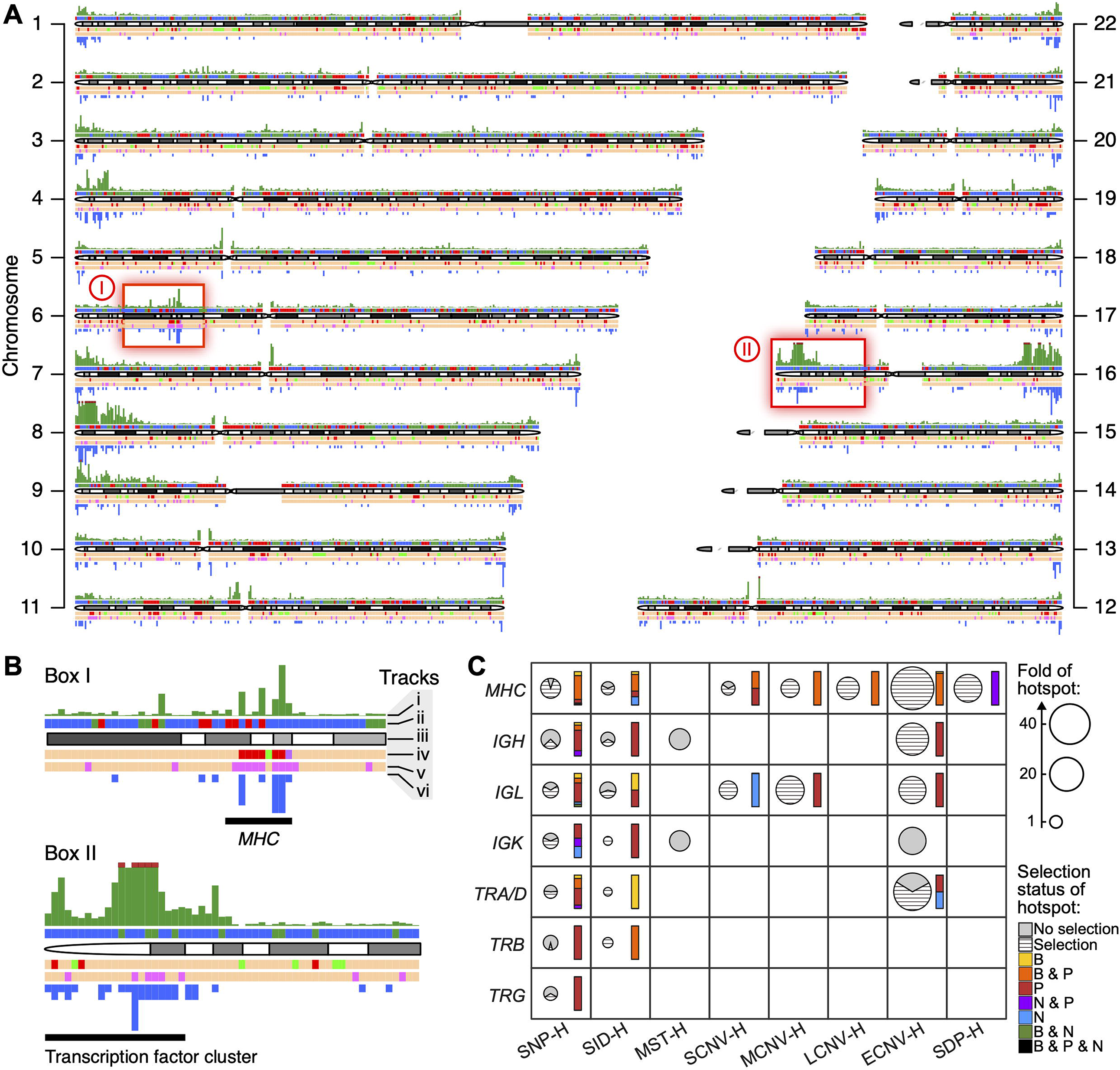
Landscape of genetic-variant hotspots and hotspot clusters. **(A)** Tracks on chromosome 1-22 show from top to bottom, in successive and non-overlapping 500-kb windows: (i) coverage of genetic-variant hotspots in window (green bar) capped at 50% by the dark red rectangles in chromosomes 8, 12 and 16; (ii) Genic zones are coloured in blue, Proximal in green, and Distal in red; ideogram; (iv) sequence windows displaying top coverage (top 10%) of positive selection hotspots (red), purifying selection hotspots (green), and both positive and purifying selections (purple); (v) sequence windows displaying top 10% coverage of balancing selection hotspots (pink); (vi) number of hotspots clusters (height of downward blue bars). Beige windows indicate where neither positive nor purifying selection hotspots reach top 10% level in track (iv), or where balancing selection hotspots do not reach top 10% level in track (v). **(B)** Tracks (i)-(vi) enlarged for red boxes I and II in (A) for the locations of *MHC* on chromosome 6, and a cluster of transcription factors on chromosome 16 [97], respectively. **(C)** Enrichment and selection-status of eight kinds of genetic-variant hotspots (x-axis) in immune system gene loci (y-axis). Fold of hotspot density inside each immunoprotein gene locus relative to autosomal average is represented by the area of the corresponding pie chart: hashed slices indicating proportion of GV hotspots overlapping, and grey slices indicating non-overlapping, with selection hotspots. Coloured bars beside each pie chart show the relative abundance of different kinds of selection hotspots acting on these GV hotspots: ‘B’ stands for GV hotspots overlapping with balancing selection hotspots, ‘P’ with positive selection hotspots, ‘N’ with negative selection hotspots. See Supplementary Table S9 for numerical data. *MHC*, major histocompatibility complex; *IGH, IGK* and *IGL*, immunoglobulin heavy, kappa and lambda loci respectively; *TRA, TRB*, *TRD*, and *TRG* are the T-cell receptor alpha, beta, delta and gamma loci. Since *TRD* is embedded in *TRA*, they are merged into a *TRA/D* locus. ‘H’ stands for ‘hotspots’.

### Formation mechanisms for hotspots and clusters

#### Contribution of recombination and retrotransposons

The association of genetic variants with homologous recombination is confirmed by the positive correlation between SNP density and recombination intensity (Figure 3A), as well as the enhancement of recombination intensity in all GV-containing windows except for the SDP-containing ones (Figure 3B). It has been proposed that retrotransposons invaded the human genome in successive waves at different times [54], and the ages of some retrotransposon subfamilies have been estimated by previous studies (Supplementary Table S5). Since the SVA, and likewise the Alu, subfamilies of similar ages show similar distribution curves in regions flanking various genetic variants (Figure 4A and Supplementary Figure S8), the SVA subfamilies are grouped together according to age to yield the SVAef (viz. SVA_E and SVA_F), SVAcd (SVA_C and SVA_D), and SVAab (SVA_A and SVA_B) subgroups; and the AluY subfamilies are grouped together to yield the ‘very young’ AluYvy (AluYa5, AluYb8, AluYb9, AluYg6, AluYf4, AluYd8, AluYa8, AluYk11, AluYh9 and AluYk12), and the ‘young’ AluYy (AluY, AluYc, AluYc3, AluYk4 and AluYf5) subgroups. LINE1 elements are likewise combined into the ‘very young’ L1vy, ‘young’ L1y, ‘middle aged’ L1m and ‘old’ L1o subgroups (Supplementary Table S5). The oldest SVAab, AluJ and L1o subgroups all show greater enrichment of SCNVs but not MCNVs, LCNVs or ECNVs in their vicinities, which is not the case with the youngest SVAef, AluYvy and L1vy subgroups (Figure 4B), suggesting that the age of these retrotransposons constitutes a significant determinant of some of the retrotransposon-GV associations. Such age effects conform to the ‘long-with-young’ (*p* = 0.018 between Younger and Older retrotransposons) but ‘short-with-all’ (*p* = 0.263) modes of association between GV length and retrotransposon age (Figure 4C). A plausible explanation is that, because the younger retrotransposons (shown in red and orange in Figure 4B) are less numerous than the older ones (shown in blue), they are more sparsely distributed in the genome. As a result, recombinations between pairs of the more sparsely distributed young retrotransposons would generate larger structural variations.

**Fig. 3.**
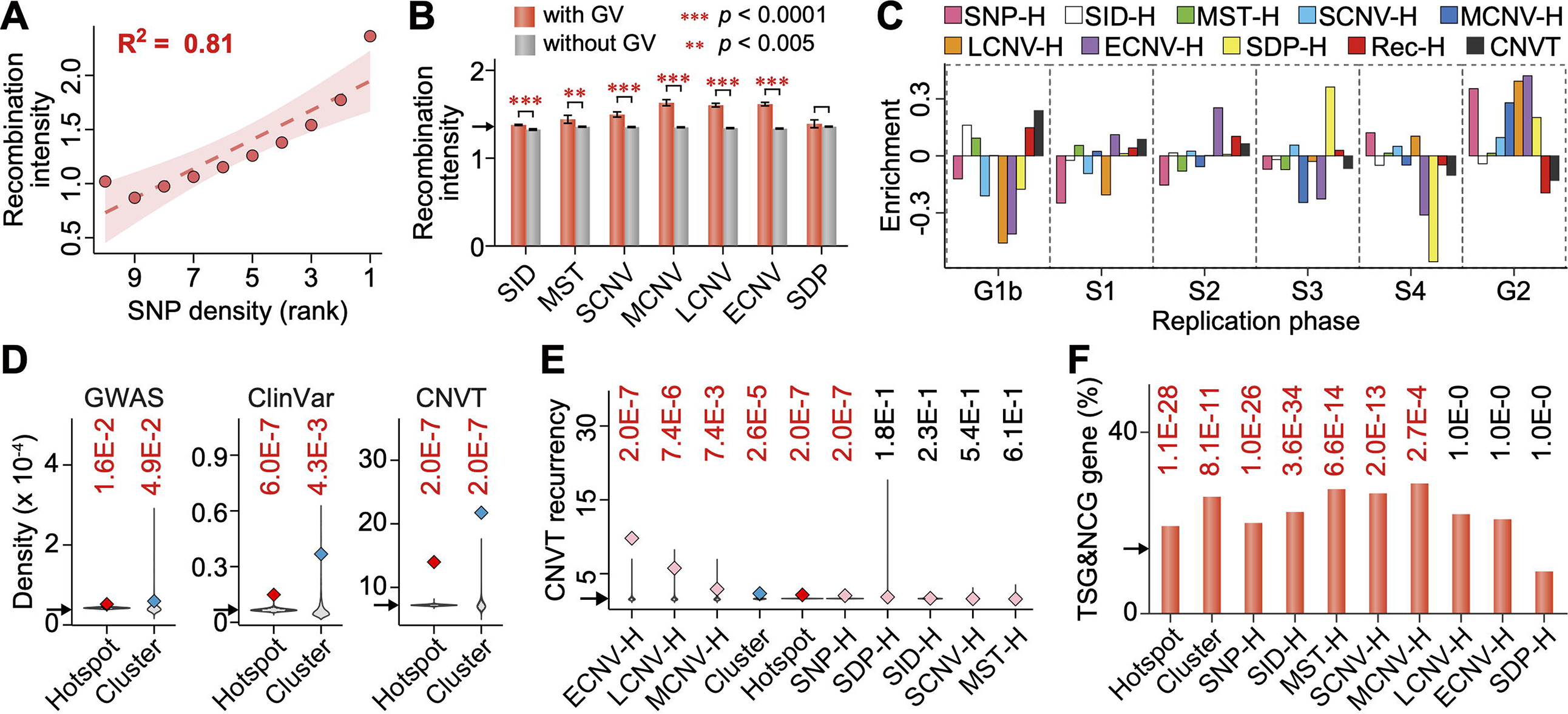
Relationships between genetic variants and genome instability. **(A)** Average recombination intensity in 10 equal groups of 1-kb sequence windows ranked according to SNP density from low (left) to high (right). R^2^ is the coefficient of determination. **(B)** Higher recombination intensity in 1-kb windows with GVs compared to windows without GVs (unpaired one-tailed t-tests, Bonferroni corrected). **(C)** Enrichment of eight kinds of genetic-variant hotspots, recombination hotspots (Rec-Hs) and somatic CNVTs in the six DNA replication phases, expressed as density fold-changes relative to autosomal average. **(D)** Densities of GWAS-identified SNPs (corrected for allele-frequency dependency, see ‘Statistical analysis’ in Methods), breakpoints of variants in ClinVar database, and somatic CNVT breakpoints in total hotspots and total clusters relative to randomly simulated genomic regions represented by the grey violin-plots. All *p*-values smaller than 0.05 are shown in red. **(E)** Recurrency of somatic CNVT breakpoints in eight kinds of hotspots, in ‘Hotspot’ representing all 44,379 genetic-variant hotspots in the genome, or in ‘Cluster’ representing all 1,135 hotspot clusters in the genome, with their *p*-values estimated by the Monte Carlo method. **(F)** Percentage of hotspot- or cluster-containing genes found in Tumor Suppressor Gene Database (TSG) and Network of Cancer Genes (NCG). The *p*-values estimated using chi-square tests are shown above each bar (Bonferroni corrected). In (B), (D) (E) and (F), arrow on y-axis indicates the autosomal average. Error bars in (B), and shaded bands around the curve in (A) indicate 95% confidence intervals. ‘H’ stands for ‘hotspots’.

**Fig. 4.**
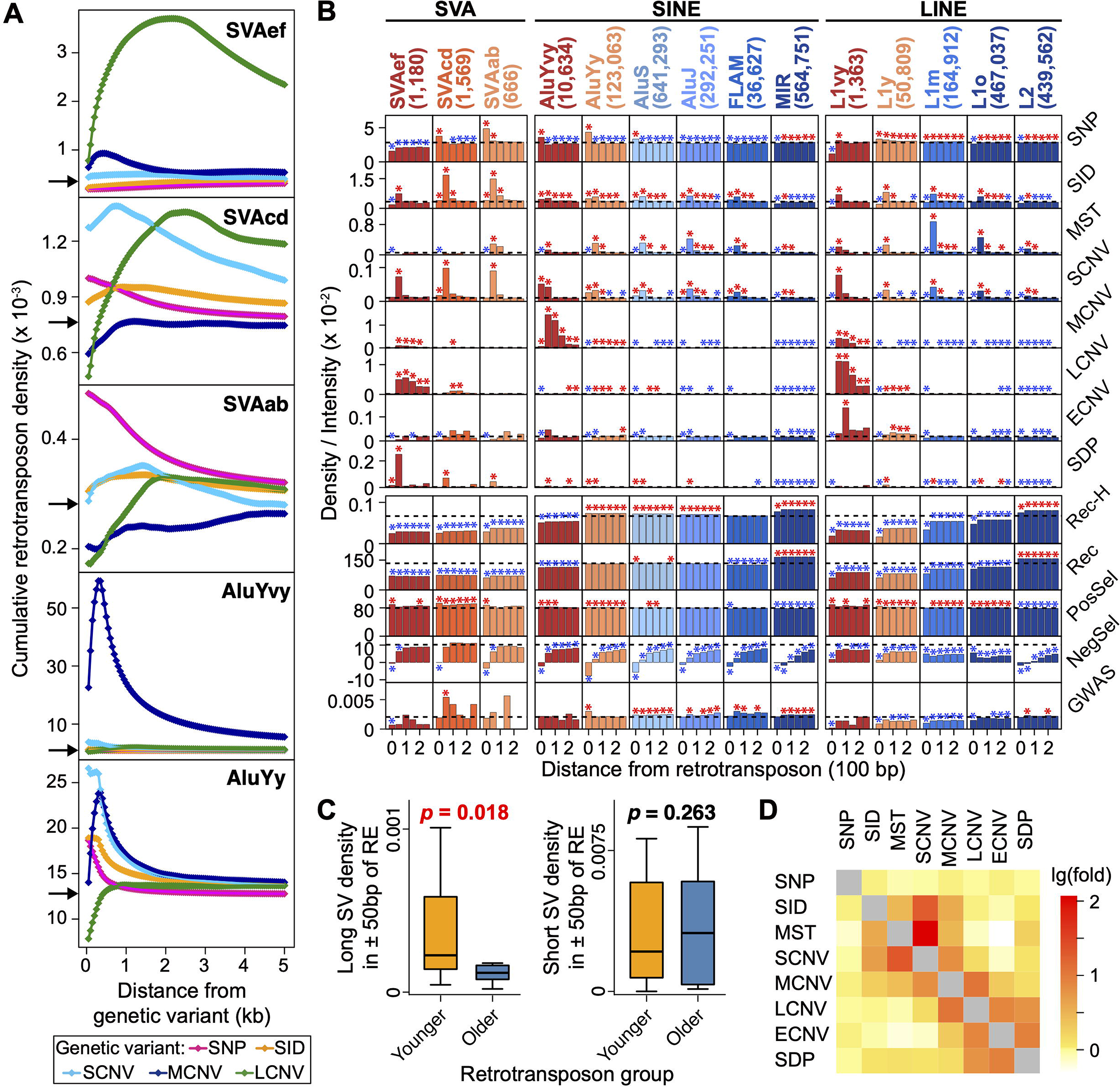
Relationships between genetic variants and retrotransposons. **(A)** Density distributions of different retrotransposon groups within (viz. at distance = 0 bp) or near SNPs, SIDs, and breakpoints of SCNVs, MCNVs and LCNVs. Arrows on y-axes indicate autosomal averages of the retrotransposon group. **(B)** Distributions of genetic variants, recombination hotspots (Rec-H), recombination intensity (Rec), positive selection (PosSel) indicated by average |nSL| values, negative selection (NegSel) indicated by phyloP scores, as well as GWAS-identified SNPs both inside (at distance = 0 bp) and within ± 250 bp of different groups of SVAs, short interspersed nuclear elements (SINE), and long interspersed nuclear elements (LINE). Designations and numbers of the retrotransposon groups indicated at the top of the columns are color-coded based on their relative ages, ranging from red for the youngest to dark blue for the oldest. Colored asterisks mark significant enrichments (red) or depletions (blue) of features based on Monte Carlo simulations (n = 1,000; * *p* < 0.005), and the dashed horizontal lines indicate the respective autosomal averages of each y-axis feature. **(C)** Density of long (MCNV, LCNV, ECNV, SDP) in the left panel, or short (SID, MST, SCNV) structural variations in the right panel, in vicinities of the younger (SVAs, AluYs, L1vy, L1y) or older (AluS, AluJ, FLAM, MIR, L1m, L1o, L2) retrotransposon groups. SV stands for structural variation; RE for retrotransposon; *p*-value < 0.05 is shown in red (unpaired one-tailed t-tests). **(D)** Enrichment of y-axis genetic variants within ± 50 bp of x-axis genetic variants expressed by the thermal scale representing the natural-log of the density fold-change relative to the autosomal average.

As a consequence of the length-age association between genetic variants and retrotransposons, the short variants (SID, MST, SCNV) form a closely co-localized group in Figure 4D, and the longer variants (MCNV, LCNV, ECNV and SDP) form a different closely co-localized group. However, although the short and long genetic variants are both associated with the young SVAef, AluYvy and L1vy in Figure 4B, the white and light-yellow squares in Figure 4D indicate that they do not show significant co-localization with one another, which suggests that other mechanisms besides retrotransposon-mediated recombination could also produce genetic variants, and natural selection may also influence the distribution of the genetic variants subsequent to their production.

#### Effects of natural selection on distribution of genetic variants

In general, the level of positive selection can be assessed based on (i) the allele-frequency spectrum, population differentiation, and (iii) haplotype structure [55]. First, using the shift of DAF to extreme values as an index of positive selection [56, 57], an elevated percentage of SNPs with DAF > 0.95, points to the enhancement of positive selection in GV hotspots and even more so in hotspot clusters relative to non-hotspot regions (Figure 5A) throughout the African, American, South Asian, European and East Asian populations (*p* < 10^−10^ for clusters in all five populations). Secondly, using population differentiation as an index, the increased values of DAF-difference between population pairs, viz. |ΔDAF| in clusters relative to non-hotspot regions, yielded *p*-values ranging from 10^−2^ down to less than 10^−13^ for eight out of ten population pairs (Figure 5B), attesting to substantial inter-population differentiation driven by directional selection within hotspot clusters during the evolution of human populations. Thirdly, using haplotype structure-based |nS_L_| statistic as an index for both soft and hard positive-selection sweeps [41], the significantly enhanced |nS_L_| scores in hotspots and even more so in clusters relative to non-hotspot regions (Figure 5C, *p* < 10^−24^ for clusters in all five populations) are likewise indicative of the presence of positive selection. A total of 26.4% of GV hotspots and 36.0% of the clusters overlap with sequence windows displaying top-5% levels in one or more of the three different measures of positive selection based on DAF, |ΔDAF| and |nS_L_| (Supplementary Table S3B), significantly more than the simulated regions (*p* = 0.0001 for both hotspots and clusters, 10,000 simulations). These three different measures of positive selection jointly enable the identification of 200,211 hotspots of positive selection based on the criterion that hotspot sequence windows display top-5% levels in at least one of the three measures (see “Determination of natural selection hotspots” in Materials and Methods).

**Fig. 5.**
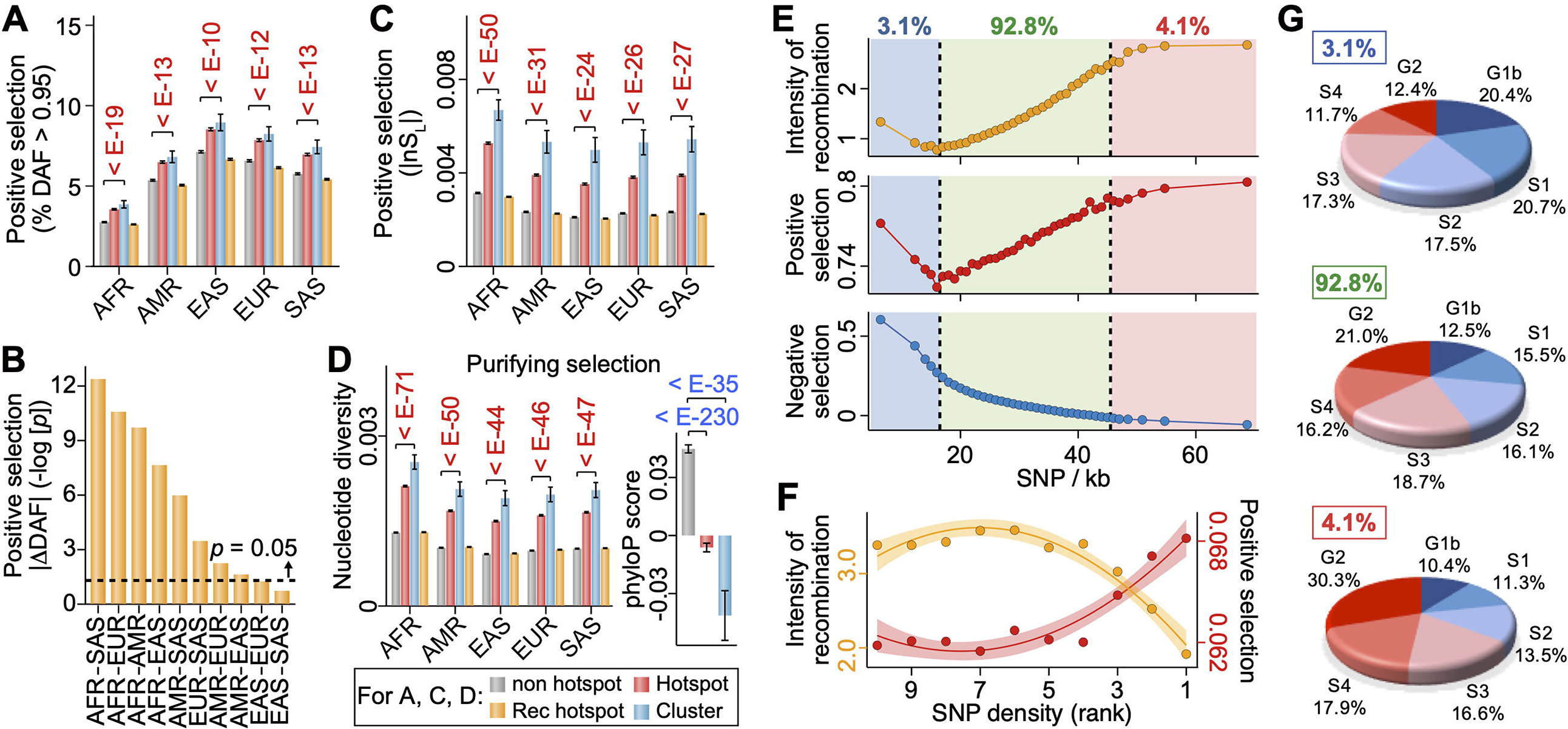
Relationships of genetic-variant density with recombination, natural selection and replication timing. **(A)** Strength of positive selection indicated by percentage of SNPs displaying DAF > 0.95 in the African (AFR), American (AMR), South Asian (SAS), European (EUR) and East Asian (EAS) populations within non-hotspot regions, total hotspots, total clusters, and Rec-Hs. **(B)** Enhancement of positive selection in clusters over non-hotspot regions in terms of |ΔDAF| between ethnic population pairs expressed by negative log_10_-transformed *p*-values from unpaired one-tailed t-tests (see ‘Statistical analysis’ in Methods). **(C)** Enhanced positive selection in clusters and hotspots relative to the non-hotspot regions indicated by elevated |nS_L_| values in five populations. **(D)** Excess of nucleotide diversity (left panel) or reduction of phyloP score (right panel) in hotspots and clusters. In (A), (C) and (D), bars share the same color-codes, and *p*-values (from unpaired one-tailed t-tests) above the bars are shown in red or in blue indicating significantly higher or lower selection in hotspots/clusters relative to the non-hotspot regions. Since no population-specific phyloP data is available, right panel in (D) compares the non-hotspot, hotspot and cluster regions of the human reference genome hg19. **(E)** Variations of recombination intensity, positive selection (in terms of average |nSL|), and purifying selection (in terms of phyloP score) in 1-kb window groups with low (3.1% of total, blue region), medium (92.8%, green region) and high (4.1%, pink region) SNP densities. **(F)** Variation of recombination intensity (orange curve) and positive selection in terms of DAF (red curve) among ten groups of SNP hotspots ranked by SNP density from low (rank 10) to high (rank 1). **(G)** Distribution of the low-, medium- and high-density windows from part (E) in the different DNA replication phases. Error bars in (A), (C), (D) and shaded bands around the curves in (F) indicate 95% confidence intervals.

Purifying selection can be assessed based on (i) reduction of genetic diversity, and (ii) cross-species sequence conservation [58]. Significant elevation of genetic diversity, measured by nucleotide diversity [59], is evident in both hotspots and clusters in all five ethnic populations (Figure 5D left panel) with *p* < 10^−44^ in all populations for the clusters. Cross-species conservation, measured by sequence conservation score measured using phyloP across 100 species [42], is significantly reduced in both hotspots and clusters relative to non-hotspot regions (Figure 5D right panel) with *p* < 10^−230^ for hotspots and 10^−35^ for clusters. Altogether 8.83% of GV hotspots, and 11.1% of the clusters, overlap with sequence windows that display either top-5% levels of phyloP score or bottom 5% of nucleotide diversity, which is significantly less frequent in comparison to simulated regions (*p* = 0 for both hotspots and clusters, 10,000 simulations). A total of 165,323 purifying or negative selection hotspots are identified as sequence windows that display at once top-5% levels of phyloP score and bottom-5% levels of nucleotide diversity.

It has been proposed that balancing selection represents as an essential adaptive force acting on structural variation hotspots harbouring genes involved in immune functions or anthropologically crucial functions [24, 25, 35]. This is confirmed by the present findings that 4.7% of the GV hotspots and 6.0% of the hotspot clusters overlap with the hotspots of balancing selection in the human genome (Supplementary Table S3B). The importance of balancing selection in shaping the distributions of hotspots and clusters is supported further by the significantly higher overlap of balancing selection hotspots with GV hotspots and the clusters relative to simulated non-hotspot regions (*p* = 0.0001 for both GV hotspots and clusters, 10,000 simulations). The findings that all three different measures of positive selection support the association of genetic-variant hotspots and clusters with positive selection (Figure 5A-C), and both of the measures of purifying selection reveal a reduction of purifying selection in hotspots and clusters relative to the non-hotspot regions (Figure 5D), therefore indicate that positive selection and balancing selection are the dominant selection forces acting on genetic-variant hotspots and clusters.

#### Co-saturation of recombination and selection in genetic variant-enriched regions

The levels of recombination and natural selection are examined at different levels of GV densities in Figure 5E. In 96.5% of the 1-kb sequence windows in the genome (light green region, Figure 5E), largely parallel variations of recombination intensity (orange curve) and positive selection (red curve) with SNP density are observed, both running expectedly opposite to the variation of negative selection (blue curve). In the windows with top 2.0% SNP densities (pink region), both the recombination and positive selection curves flatten as they approach saturation. This points to the possible underestimation of recombination rate in the presence of strong positive selection and vice versa. Mechanistically, excessive sequence shuffling by recombination events would limit the effectiveness of positive selection in terms of the attainable DAF, leading to the low percentage of SNPs with high DAF as well as low |nS_L_| in over 30,000 previously determined recombination hotspots (Rec-Hs) in the human genome [60, 61] of all the ethnic groups (orange bars, Figure 5A and C). On the other hand, strong positive selection could eliminate some of the unfit recombinant genotypes, thereby diminishing the footprints of the recombinations and causing an underestimation of their effects. On this basis, the densities of genetic variants, being independent of allele or haplotype frequencies, could represent a more robust measure of genetic diversity than frequency-dependent measures where selection and recombination are highly active. Interestingly, in the SNP hotspots that are 1^st^ or 2^nd^-ranked in terms of SNP density (Figure 5F), recombination decreases yet the level of positive selection remains elevated, suggesting that although recombination tends to be underestimated owing to the presence of positive selection, the positive selection is too strong to be suppressed by recombination. Among different GV hotspots, those of SNP, ECNV and LCNV show the largest percentage of overlap with positive selection hotspots (Supplementary Table S3B), pointing to the presence of exceptional positive selection within these three kinds of hotspots.

In the windows with the lowest-1.5% SNP densities (light blue region, Figure 5E), low SNP is accompanied by high recombination intensity and positive selection. This finds a possible explanation in the top pie chart of Figure 5G, which shows that over 50% of these lowest-1.5% SNP-density windows are in fact replicated in the G1b-S2 phases where the fidelity of DNA replication is high [62–64], suppressing thereby the occurrence of SNPs (see ‘Replication-time segments’ in Materials and Methods for the classifications of sequence windows into six replication phases).

### Impacts of hotspots and clusters on function and disease

#### Idiosyncratic association with functional genomic features

It has been observed previously that Alu insertions of different ages enhance SNP occurrences in their neighborhoods differentially [29], suggesting that genomic feature-genetic variant associations vary with the nature of the genetic variant or the genomic feature or both. Accordingly, the co-localizations of eight kinds of GV hotspots with a wide spectrum of genomic features are analysed in Figure 6. Some of the co-localizations turn out to be highly idiosyncratic, exemplified by the strong associations of the DNA methylation features with the SNP hotspots; or the large intergenic non-coding RNAs (LINC) with ECNV and SDP hotspots; and histone modification features with MCNV hotspots.

**Fig. 6.**
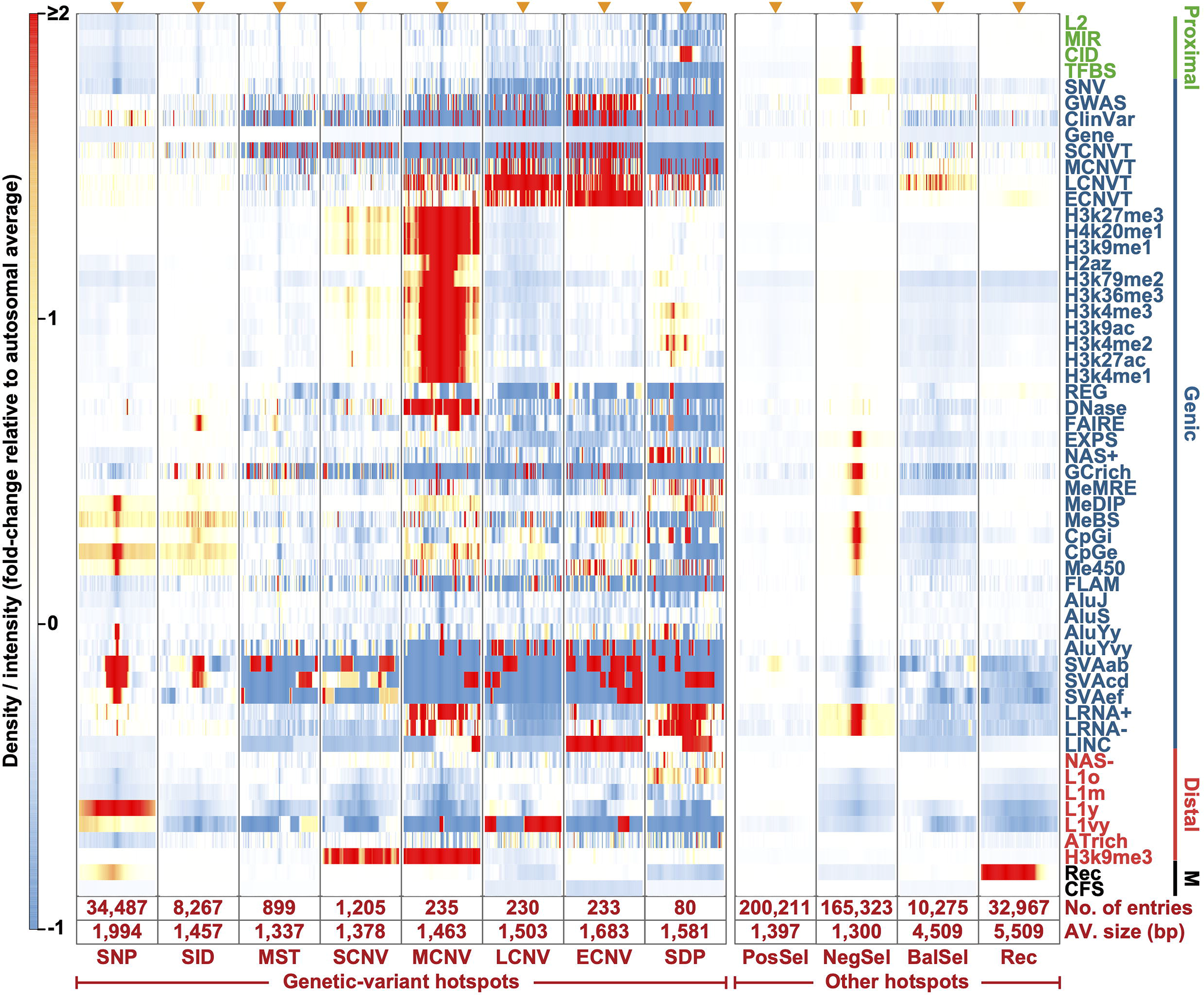
Genomic features inside and near genetic-variant, selection and recombination hotspots. The density or intensity of 55 types of genomic features occurring between 4-kb upstream (left end) to 4-kb downstream (right end) from the centre (indicated by orange arrowhead at the top of each column) of different kinds of hotspots, expressed in fold-changes relative to the respective autosomal average in accordance to the red-blue thermal scale. The number and average size of different kinds of hotspots are shown in the bottom two rows of the figure. The genomic features indicated on the y-axis are grouped into Proximal-zone (green), Genic-zone (blue), and Distal-zone (red) features based on their co-localization relationships (Supplementary Figure S2) [35], along with Marker (M) features (black). The descriptions and abbreviations of different genomic features are given in Supplementary Table S1A.

The dissimilar properties of the four size-classes of CNVs are confirmed by the strong associations of histone-modification features such as H3k9me3 with the hotspots of MCNV and SCNV but not those of LCNV and ECNV; the much stronger associations of H2az, H3k79me2 and open chromatin elements (DNase) with hotspots of MCNV relative to those of SCNV; and the strong association of LINC with the hotspots of ECNV relative to those of MCNV, or even more so those of LCNV or SCNV (Figure 6 and Supplementary Table S1A). The PosSel-Hs co-localize mildly with the SVAab retrotransposons. The NegSel-Hs co-localize positively with a range of regulatory features, methylation sites and disease-related sites, while their negative co-localizations with the Alu, SVAab, SVAcd and L1 retrotransposons support the suggested paucity of purifying selection in transposable elements [65, 66].

The 1,135 hotspot clusters in the human genome comprised twenty-three different combinations of hotspots, a large majority of which include the hotspots of SNP and hotspots of structural variants. Each kind of hotspot clusters is associated with its own characteristic array of genomic features. The associations of for example the long RNA (LRNA-) with the SNP+MCNV cluster (viz. clusters comprising the SNP and MCNV hotspot) are largely additive, not far from the sum of the individual associations of LRNA-with SNP hotspot and MCNV hotspot; in contrast, the associations of H3k27ac toward the SNP+SCNV cluster are synergistic, far exceeding the sum of the individual associations of H3k27ac with SNP hotspot and SCNV hotspot (Supplementary Figure S9).

#### Distributions at immune system gene loci

Although genes are not specially co-localized with GV hotspots or clusters (Figure 6 and Supplementary Figure S9), the effects of genetic variants and natural selection are often discernable within particular gene clusters (Figure 2B), as illustrated by the immune system gene loci including *MHC* where protein sequence diversity represents a necessity [21]. Thus the immunoglobulin *IGH*, *IGL*, and *IGK* loci, and the *MHC* locus all overlap with clusters that are enriched with various kinds of hotspots, such as a 40-fold enrichment of ECNV hotspots in the *MHC*-locus relative to genomic average (Figure 2C). The GV hotspots also exhibit abundant co-occurrence of positive in terms of PosSel-H and balancing selection in terms of BalSel-H contents (orange bars, Figure 2C), but little purifying selection in terms of NegSel-H content (blue bars). These results suggest that the high sequence diversity of the *IG*- and *MHC*-loci has been achieved mainly through positive and balancing selection within the hotspots and clusters. The weaker presence of GV hotspots as indicated by reduced pie size, or selective processes as indicated by hashed slices, in Figure 2C among the *TRA*, *TRB, TRD* and *TRG* loci of T-cell receptors are consistent with the reduced need by these receptors for sequence diversity compared to the *IG*- and *MHC*-loci.

#### Functional evolution in late-replicating DNA

A well-defined temporal order of DNA replication is characteristic of normal cell division. The periods of genomic DNA synthesis have been classified, from early to late, into the G1b, S1, S2, S3, S4 and G2 phases in fifteen cell types [46], thus allowing the partition of 1-kb autosomal windows into these six phases (see “Replication-time segments” in Materials and Methods). Genic zones are enriched in the early-replicating G1b, S1 and S2 phases, Distal zones in the late-replicating S4 and G2 phases, and Proximal zones in the intermediate S3 and late-replicating S4 phases (Figure 1C). The density of recombination hotspots declines from G1b toward G2 (red bars, Figure 3C), in accord with the previous finding of a reduction of homologous recombination in late-replicating DNA [33]. High frequencies of GV hotspots are found in G2-phase DNA despite a low level of recombination hotspots.

The DNA sequence windows that undergo early replication in phase G1b are enriched in numerous functional genomic features including the histone-modification sites and gene/regulatory sites throughout the Genic, Proximal and Distal zones (red rectangles, Figure 7A top panel), whereas sequence windows in the late-replicating phases S3-G2 are relatively depleted of these functional features (blue rectangles). Nonetheless, within these function-depleted phases, the presence of functional features is more evident in regions where hotspots and clusters locate, especially the histone-modification sites in the Distal-zone hotspots and clusters (red rectangles in middle and bottom panels, Figure 7A). Six groups of gene pathways are found enriched in the Distal zones (Figure 7B and Supplementary Figure S10). Among them, the autoimmune thyroid disease, antimicrobial humoral response, natural killer cell, xenobiotic metabolism, epidermal protein disulfide-binding, sensory perception, and neuroactive ligand-receptor interaction pathways are all associated with GV hotspots (red and pink circles, Figure 7B). Moreover, the latter two pathway groups display the foremost hotspot enrichment, with 46% and 64% of their genes respectively (black slices, Figure 7C) showing the co-existence of positive, balancing and purifying selections, in contrast to the immune system gene loci which are dominated by positive and balancing selections (Figure 2C). These findings indicate that the presence of hotspots and clusters could attract various kinds of natural selection forces that can facilitate pathway development even in regions of the genome that usually harbor relatively few functional genomic features.

**Fig. 7.**
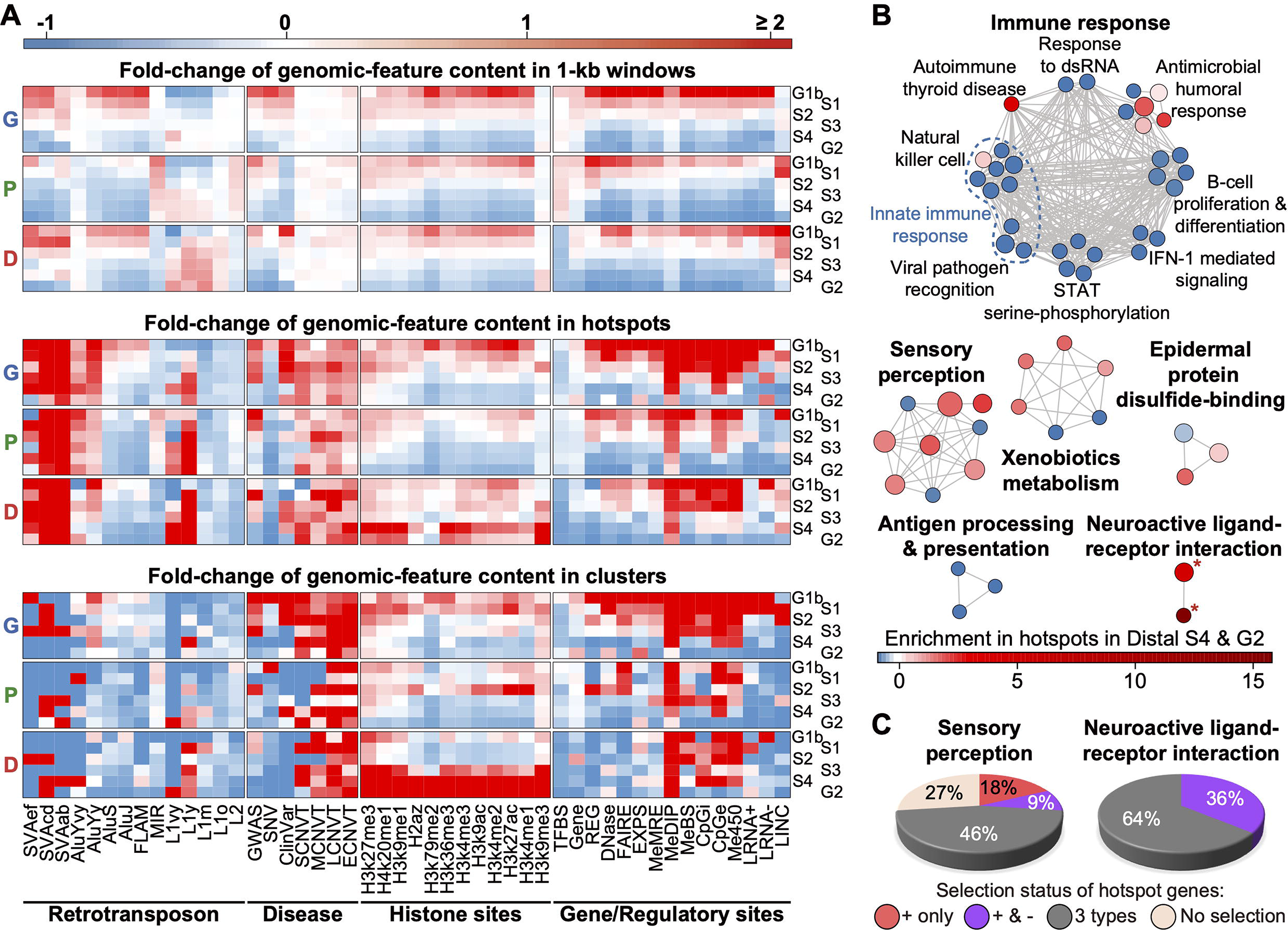
Distributions of genomic features in different replication phases and genomic zones. **(A)** Levels of various genomic features (as shown on x-axis) within all 1-kb windows (upper panel), all hotspots (middle panel), or all clusters (lower panel) belonging to replication phases G1b-G2 in the Genic (G), Proximal (P) or Distal (D) zones, in accordance to the red-blue thermal scale for fold-change over autosomal average (see Supplementary Table S10 for data on genomic-feature enrichments). **(B)** Enrichment Map for Distal-zone genes annotated using g:Profiler [98] based on Gene Ontology biological process [99] and KEGG [100]. Each node represents a pathway (with 3 to 350 genes) significantly enriched in Distal zones with Benjamini-Hochberg false discovery rate < 0.05, and node size is proportional to the number of Distal-zone genes belonging to the pathway (see pathway IDs in Supplementary Figure S10). Pathways are connected by a grey edge when they share ≥ 50% genes. Node color, represented by the red-blue thermal scale, indicates the fold-change of the fraction of the genes over the fraction of 1-kb sequence windows that overlap with hotspots in the Distal-zone S4 and G2 phase DNA (see Supplementary Table S11). Asterisks mark pathways that overlap with cluster(s). **(C)** Selection status of pathway genes that overlap with hotspots in the Distal-zone S4 and G2 phases. Fractions of genes overlap only with PosSel-Hs (red slices), with both PosSel-Hs and NegSel-Hs (purple slices), and with all (black slices) or none (beige slices) of PosSel-Hs, NegSel-Hs and BalSel-Hs.

#### Enrichment of disease-related variants in hotspots and clusters

The close relationships of genetic-variant hotspots and clusters with different functional genomic features (Figure 6 and Supplementary Figure S9) suggest that perturbations in the DNA sequences in these regions may result readily in disease-related mutations. In accord with such expectation, the densities of germline SNPs identified from GWAS, the breakpoints of disease-related germline variants in the ClinVar database, and the breakpoints of somatic tumor CNVTs are all significantly enriched within the GV hotspots and clusters relative to simulated regions in the human genome (Figure 3D). The density of GWAS-identified SNPs within the total GV hotspots is increased to 1.4-fold, to as high as 7.1-fold in ECNV hotspots, or synergistically to 19.0-fold in the SNP+ECNV clusters relative to the autosomal average, far exceeding the sum of their separate enhancements in the SNP and ECNV hotspots (Supplementary Table S6). Instead, clusters formed by SNP and SID does not have this kind of synergistical elevation, but rather a reduced level of disease-associated variants relative to their separate hotspots. This observation suggests that the potential role of SNP+ECNV clusters are pivotal genomic regions attributable to complex traits and diseases. Furthermore, the GWAS-identified SNPs in the GV hotspots or the clusters are characterized by comparable risk allele frequencies, but higher effect sizes and contributions to heritability, relative to the GWAS-identified SNPs in non-hotspot regions (Supplementary Figure S11). Besides, the GWAS-identified SNPs are found significantly enriched inside or near SVAcd, AluYy, AluS, AluJ, FLAM, MIR and L2 retrotransposons but not the youngest SVAef, AluYvy and L1vy retrotransposons (Figure 4B).

The density of CNVTs is likewise increased to 1.9- and 2.9-fold within total hotspots and clusters respectively; or as high as 13.7-fold in the LCNV hotspots and 28.3-fold in the ECNV hotspots (Supplementary Table S6), demonstrating thereby the association of hotspots and clusters with enhanced germline as well as somatic disease-related mutations. Upon division of the somatic CNVTs into the four size-classes of SCNVT, MCNVT, LCNVT and ECNVT as in the case of the germline CNVs, LCNVTs are most strongly associated with LCNV hotspots, and ECNVTs most strongly with ECNV hotspots (Figure 6), indicating that LCNV hotspots represent significant sites of LCNVT formation, and ECNV hotspots represent significant sites of ECNVT formation, in keeping with the earlier finding that recurrent germline CNVs provide a useful basis for predicting cancer susceptibility [67]. CNVT recurrency in tumours is particularly strong in the ECNV, LCNV, and MCNV hotspots, in clusters, and in SNP hotspots (Figure 3E). Such enrichments of disease-related variants in the hotspots and clusters are observed regardless of the genomic zone or replication timing of the DNA sequences (Figure 7A middle and bottom panels).

There are in the human genome 448 genes that contain one or more clusters, and 24 genes that contain more than two clusters, reaching a maximum of nine clusters in the diabetes-related *PTPRN2* and the potential tumor suppressor *CSMD1* (Supplementary Table S4). Within 33 of these cluster-containing genes, the cluster portions are significantly enriched with somatic CNVTs compared to non-cluster portions (see “Somatic CNV breakpoints in cluster-containing genes” in Methods), which attests to the extraordinary effectiveness of clusters in the generation of CNVT mutations. Among the 448 cluster-containing genes, 113 genes are found in the Network of Cancer Genes 6.0 [68] or the Tumor Suppressor Gene Database 2.0 [69], revealing significant overlaps between cancer genes with hotspot clusters (*p* = 8.1 × 10^−11^, Figure 3F), or with hotspots of SNP, SID, MST, SCNV or MCNV (Figure 3F).

## Discussion

### Formation mechanisms for common and rare variants in hotspots and clusters

SNPs and structural variations in GWAS are widely employed as common and rare genetic markers respectively the “Common Disease-Common Variant’ and ‘Common Disease-Rare Variant’ research strategies [5, 12, 13]. In the present study, they are both found to be concentrated in hotspots and clusters that cannot be formed due to background mutations, in a clear departure from the neutral evolution model [70–73]. The presence of numerous hotspot clusters that comprise an SNP hotspot along with the hotspot of some structural variations (Figure 2 and Supplementary Figure S9) suggests that the formation mechanisms for the SNP hotspots overlap with those for the hotspots of SIDs, MSTs, SDPs and CNVs of different lengths in the same clusters. It follows that the different structural variations do exhibit overlaps among themselves, as exemplified by the co-localizations between copy number gains and copy number losses, insertions and tandem duplications, as well as duplications and SDPs, supporting that the unstable genomic regions are prone to diversified genetic variants (Supplementary Figure S12). Evidence suggests that homologous recombination can be mutagenic, producing SNPs due to the error-prone nature of the DNA polymerase involved, or generating structural variations via ectopic homologous recombination events between repeated sequences [30, 32, 74, 75]. As well, the association of homologous recombination with the formation of a range of genetic variants is indicated by the co-occurrence of elevated genetic-variant density and high recombination intensity (Figure 3A and B). However, only 15.0% of GV hotspots overlap with the previously identified recombination hotspots (Supplementary Table S7), possibly because of underestimation of recombinations on account of the reduction of their footprints by the strong positive selection in the GV hotspots (Figure 5F), or by other mutagenic mechanisms such as replication-based mechanisms or the non-homologous end joining that contributed to the formation of the GV hotspots and clusters. This is demonstrated by the high concentrations of GV hotspots in G2 phase-replicated DNA despite the relative scarcity of recombination hotspots in such DNA (Figure 3C), coinciding with the stronger presence of non-homologous end joining compare to homologous recombination in G2-phase DNA [33].

It is noteworthy that nucleotide differences between highly similar sequences (e.g. SDPs) located at different genomic positions may be falsely identified as SNPs due to mismapping of short-reads [25], giving rise to biased associations of SNPs with structural variations. On this basis, false-positive SNPs may be expected to be particularly frequent near SDPs where the repetitive sequences would be conducive to such mismapping. Since SNP densities are much lower in the vicinity of SDPs relative to the vicinities of MCNV, LCNV and ECNV (Supplementary Figure S12), subtraction of the level detected in the vicinity of SDP from the SNP levels observed from the vicinity of the various CNVs may be considered as a useful correction measure for those SNPs that are neat the MCNV, LCNV and ECNV.

### Negative effects of hotspots and clusters

The suggestion that retrotransposons could mediate homologous recombination [26, 27] is confirmed by the close co-localizations of a range of genetic-variant hotspots and clusters with different retrotransposons, and the positive co-localization of AluYy insertions not only with various hotspots and clusters but also with recombination features (Figures 4B and 6). Such findings indicate that GV hotspots and clusters harbor sites of genomic instability which expedite the production of neutral, advantageous or detrimental variants through recombination. The detrimental effects are evident in the strong associations of hotspots and clusters with cancerous CNVTs, and the enrichment of disease-associated variants in GWAS (Figure 3D-F). Furthermore, the parallel declines in Rec-H and CNVT in G1b- and G2-replicating DNA (Figure 3C) support the proposal of homologous recombination as a source of CNVTs [74]; and the strong associations between LCNVT and LCNV hotspots, as well as between ECNVT and ECNV hotspots (Figure 6) suggest that the selfsame sets of cis-acting retrotransposons are utilized as nucleation points for the formation of both germline CNV and somatic CNVT through recombination [76, 77]. Moreover, clusters are also observed for somatic structural variations implying mechanistical linkages of genetic variants in those clusters [78]. However, the germline CNVs and somatic CNVTs are dissimilar in their length distributions, resulting in a merger of the four distinct peaks of germline CNVs into a more even distribution of CNVTs of different lengths (Figure 1A, bottom panel), as well as in a more parallel distribution of CNVTs compared to germline CNVs with Rec-H in DNA sequences belonging to different replication phases (Figure 3C). These dissimilarities between CNVTs and germline CNVs could have arisen from the prolonged exposure of the latter to natural selection. The association of CNVTs with GV clusters is particularly strong: there are 33 genes in the genome where CNVT breakpoints are significantly enriched within segments bearing GV clusters relative to other segments of the gene (overall Bonferroni-corrected *p*-value < 0.05). The significance of such enrichment reaches *p* < 10^−58^ for *CACNA1C* and *SNX29* from the Network of Cancer Genes, and *p* < 10^−61^ for *WWOX* and *CSMD1* from the Tumor Suppressor Gene Database (Supplementary Table S4). Therefore the cluster segments of such genes clearly rank among the hotspots of oncogenesis within the genome.

### Positive effects of hotspots and clusters

GV hotspots and clusters produce advantageous genetic variants that could be further elevated in frequency by positive selection to expedite functional adaptation. This is illustrated by: (i) enrichment of SNPs with DAF > 0.95 in hotspots and clusters in all five ethnic populations (*p* < 10^−10^, Figure 5A), accompanied by large overlaps of PosSel-Hs with SNP, LCNV and ECNV hotspots (Supplementary Table S3B); (ii) rise of the inter-population differentiation |ΔDAF| owing to positive selection within the clusters (Figure 5B); and (iii) increase of haplotype homozygosity, measured by |nS_L_|, on account of positive selection in all five ethnic populations (*p* < 10^−24^, Figure 5C). Outstanding examples are provided by the *MHC-* and *IG*-loci which are enriched with SNP, SID, MCNV, LCNV, and ECNV hotspots associated with positive and balancing selections (Figure 2C), revealing the exceptional hotspot- and cluster-driven evolutionary development of these immune system loci.

Another example of the positive functional effects of hotspots and clusters is provided by the development of the olfactory and taste sensory pathways, and neuroactive ligand-receptor interaction pathways through collaboration of balancing, positive and purifying selections in the hotspot and cluster regions that amount to only 3.3% of the total base pairs in these otherwise functional deserts of the S4- and G2-replicating Distal zones (Supplementary Table S8), which clearly demonstrates the important advantage of hotspots and clusters as incubators for the accelerated adaptive evolution. The dominance of purifying selection in the protein-coding regions of genes [79] suggests that a majority of the coding sequences in the human genome are already largely optimized by positive selection, and therefore in need of shielding by purifying selection against adverse sequence alterations. On the other hand, the much weaker purifying selection than balancing or positive selection observed in the immunoprotein genes is consistent with the younger age of these genes, where evolutionary development might still be ongoing, propelled by genetic variants working in conjunction with balancing and positive selections.

The coincidence between hotspots of GVs and positive selection may seem paradoxical, for one might suppose that the selection process would sweep away the GVs. However, a high level of the nSL statistic could serve as an indicator for the existence of ‘soft sweep’ where genetic variants or multiple variants at a single locus remain standing until the sum of their allele frequencies reaches one [41]. Genetic forces and evolutionary history such as the complex demographic history of human populations may render difficult a rigorous description of positive selection by any single measurement [56, 80]. For example, population bottlenecks can lead to the shift of DAF to extreme values; a population expansion may increase ΔDAF between population pairs; and the nSL statistic may vary with the demographic model employed. Therefore, the observation of enhancement of positive selection in GV hotspots and clusters based on all three measures for detection of positive selection signals improves the reliability of the observed enhancement.

### Missing heritability due to co-occurrence of recombination and positive selection

Genetic components associated with complex traits and disorders could be missed in genomic regions simultaneously subjected to high levels of positive selection and recombination. One of such cases has been well evident in a 3,551-bp segment of the schizophrenia-associated *GABRB2* gene that codes for the β2 subunit of GABA_A_ receptor. The derived alleles of a number of SNPs within this segment are under recent and ongoing positive selection, for they increase through alternative splicing the production of the longer isoform of the β2 subunit and diminish the production of the shorter β2 subunit favoured by the ancient alleles [81]. The ancient alleles become less fit than the derived alleles under modern conditions, resulting in positive selection of the derived alleles. The same has been observed in some other genes [82–85]. In the meanwhile, this segment harbours an AluYi6 insertion, which serves as a recombination centre to largely enhance recombination rate within the segment [86]. Under such circumstances, the elevated recombination can play positive and negative roles simultaneously. On the one hand, novel sequence variations and haplotypes are brought about by recombination for positive selection to increase the frequencies of the advantageous genotypes. On the other hand, sequence alternations could also lead to functional perturbations that associate with the etiological basis of schizophrenia [87]. The co-occurrence of recombination and positive selection points out a possible genetics mechanism for the development of complex diseases.

A wide range of schizophrenia-like phenotypes displayed by the *GABRB2*-knockout mice, and the reversal of these phenotypic alterations by antipsychotic drug further underline the pivotal role of *GABRB2* in the development of schizophrenia [88]. As well, the association between *GABRB2* and schizophrenia has been validated by multiple genetic studies on different ethnic populations [89–91], and SNPs, haplotypes and CNVs in the 3,551-bp segment are found to be associated with schizophrenia taking a candidate gene approach [92, 93]. However, genome-wide association studies report no significant association between any of the SNPs in this 3,551-bp segment and schizophrenia. Fine-resolution linkage disequilibrium analysis in this segment reveals much higher recombination in the ancestral allele-containing haplotype groups relative to the derived allele-containing haplotype groups, suggesting active recombination in this region together with intense positive selection on the derived alleles [86]. Such complexity due to the co-occurrence of recombination and positive selection, interacting with each other to affect local genetic diversity landscape, could blur the allele and haplotype signals thereby potentially contributing to the missing heritability problem in GWAS. The depletion of GWAS-identified SNPs near the youngest (viz. SVAef, AluYvy and L1vy), versus enrichment near the older (viz. AluS, AluJ, MIR etc.), retrotransposons might point to missing associations stemming from the elevated levels of recombination and positive selection in the youngest retrotransposons (Figure 4B). Therefore, the genomic regions that are subject to pronounced recombination and positive selection, such as the genetic-variant hotspots and clusters or the vicinity of young retrotransposons, would merit close investigation, more reliance also may be accorded to genetic variants, such as CNVs, that are more resistant to the obscuring effects of recombination together with selection.

### ‘Common Disease-Hotspot Variant’ hypothesis

The dissimilar associations of SCNV and MCNV hotspots with histone-modification sites, SNP and SID hotspots with methylation-related sites, and LCNV and ECNV hotspots with somatic CNVTs (Figure 6) are indicative of the specialized deployment of different genetic variants in the genome to meet specific functional needs. Notably, the associations observed between hotspots and genomic features are unlikely stemmed from the similarity in their detection methods, because data on different genomic features and genetic variants were generated by independent projects with different experimental methods. The strength of association of GV hotspots and clusters with function is underlined by the findings that 34.8% of total GV hotspots and 43.9% of total clusters display some balancing, positive or negative selection. As a result, sequence perturbations in these pivotal sites may be expected to give rise to phenotypical changes including human disorders. It is noteworthy that searches for phenotype-genotype associations are usefully guided by the ‘Common Disease-Common Variant’ and ‘Common Disease-Rare Variant’ strategies. However, the former is limited by the fraction of heritability of complex traits explained by common SNPs, which is highly variable (20-90%) for different traits and different studies [94, 95], and the latter is confronted by the difficulty of finding the rare variants [12, 35], both leaving substantial proportions of heritability unexplained. The ‘hypothesis driven’ strategies have been found useful in certain cases, but they usually require some measure of prior knowledge about the disease mechanism. It has also been found that incorporation of functional genomic data in association studies also can increases the discovery of trait/disease-associated sites by about 5% [96]. As well, it is important that the full implications of the sequence changes also should not be missed. In our study, for example, the medium size MCNVs are correlated to histone modification sites, as indicated in Figure 6. Accordingly, when an investigator finds that some disease displays correlation with an MCNV alteration, he would be encouraged to investigate in depth any histone modification sites in the vicinity, thereby reducing the chances of missing some important functional implication of the MCNV observed. Accordingly, in view of the evident elevations of GWAS discovery rates in GV hotspots and clusters, and the enriched presence of both LCNVTs in LCNV hotspots and ECNVTs in ECNV hotspots, intensive efforts directed to the discovery of common or rare genetic variants in hotspots and clusters, and characterization of their potential association with complex disease and traits, appear warranted in order to search for some of the missing heritability.

## Conclusion

Overall, in the present study, a high-resolution examination has been conducted throughout the Genic, Proximal and Distal genomic zones in the human genome on eight kinds of germline genetic variants retrieved from public databases. Identification of genetic-variant hotspots and clusters based on regional densities of genetic variants, independent of allele and haplotype frequencies, minimizes underestimation on account of positive selection and recombination saturation. The findings on their distributions, formation mechanisms, associated genomic features, as well as their effects on functional development, complex diseases and traits underline the pervasive and double-edged sword nature of their impact on the human genome (Figure 8). They advance functional genome evolution, but they also constitute important foci of cancers and other diseases, both arising from their association with genome instability. The usages of common and rare variants to search for disease-GV associations, as formulated in the ‘Common Disease-Common Variant’ and ‘Common Disease-Rare Variant’ hypotheses respectively, have brought about important advances in the understanding and detection of the genomic changes related to phenotypical changes, it is recognized that they yet face the problem of missing heritability, where the genomic sites associated with diseases and traits could only be partly located. In view of this, and the enrichment of both common and rare genetic variants in GV hotspots and clusters, it is proposed that ‘Common Disease-Hotspot Variant’ hypothesis could provide an additional, complementary approach. The basis of this hypothesis is that common diseases and traits are frequently attributable to genetic variants, common as well as rare types, occurring in regions marked by extraordinary genome instability. Such instability at these unstable regions would enhance the probability of not only germline variants, but also disease- or trait-associated germline variants and disease-related somatic variants. Thus, the GV hotspots and clusters represent an unexpected source of disease-related heritability that could easily be missed due to the effects of local recombination at or near saturation levels along with directional selection on allele or haplotype frequencies, and intensified discovery and mutation monitoring of hotspots and clusters could help to uncover some of this missing heritability.

**Fig. 8.**
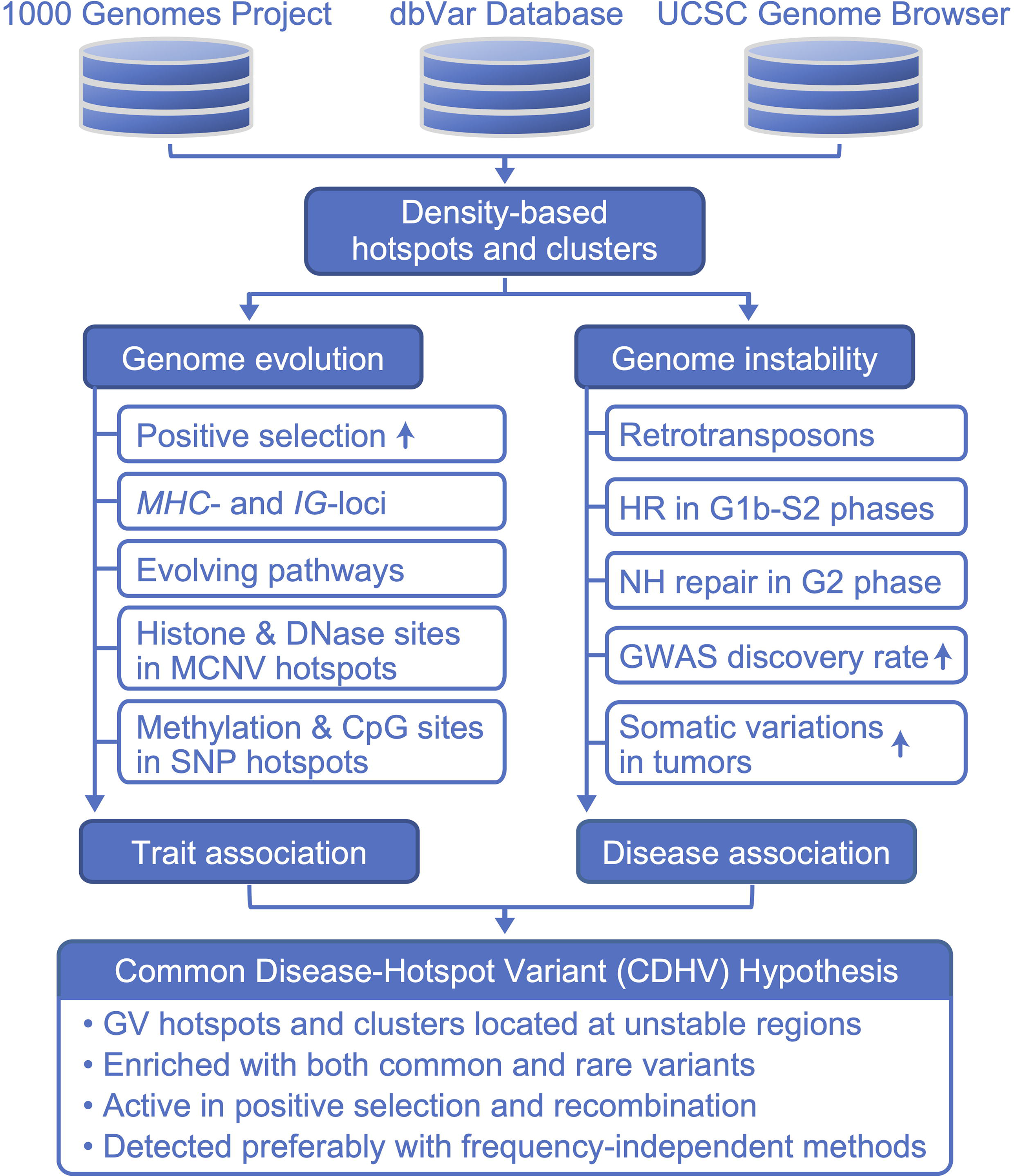
Schematic summary of present findings. Public genetic-variant databases are employed to identify density-based hotspots and clusters, which are found to be dynamic centres with accentuated positive and/or negative changes in the genome. On the one hand, they work in conjunction with positive selection to enhance sequence diversity, evolution of specific genes and pathways, and development of functional genomic features. On the other hand, they are co-localized with destabilizing retrotransposon elements, and occurrences of homologous recombination (HR) in the early DNA replication phases, and non-homologous (NH) repair in the late DNA replication phases. Their associations with high GWAS discovery rates and somatic variants signal the importance of associations with complex traits and diseases, and therefore their utility as the basis for a potential ‘Common Disease-Hotspot Variant’ strategy in a search for the missing heritability in association studies.

Since the landscape of the unstable regions may be complicated by saturation of recombination and selection, a Common Disease-Hotspot Variant hypothesis-based approach requires experimental and statistical methods that are relatively insensitive to recombination and selection. In this regard, structural variations such as CNVs will serve as valuable markers, for they are, compared to SNPs less prone to mutation saturation arising from increased recombination. GV density-based statistical methods are also advantageous on account of their relative immunity to directional selection as compared to statistical methods of population genetics commonly used in association studies, which are often frequency-based and susceptible to the effects of selection on allelic and haplotype frequencies. Therefore, through additional focus on GV hotspots and clusters, and increased adoption of frequency-independent statistical methods, some of the missing heritability may come to be captured, deepening insight into the genetics of complex disorders and traits, and improving the detection of disease- or trait-associated variants.

## Supporting information

Supplementary Dataset S1. Genomic coordinates of all hotspots identified in the present study.

Supplementary File S1. Supplementary figures S1-S12, captions of Supplementary Tables S1-S11, and description of Supplementary Dataset S1.

## List of Abbreviations

SNP: single-nucleotide-polymorphism
CNV: copy-number-variation
GV: Genetic variant
GWAS: genome-wide association study
SID: small indel
MST: microsatellite
SDP: segmental duplication
PosSel-H: positive selection hotspot
NegSel-H: negative selection hotspot
BalSel-H: balancing selection hotspot
Rec-H: recombination hotspot
DAF: derived-allele frequency
MAF: minor-allele frequencies
LINC: large intergenic non-coding RNAs
DNase: open chromatin elements
LRNA-: long RNAs

## Declarations

### Ethics approval and consent to participate

Not applicable.

### Consent for publication

Not applicable.

### Availability of data and materials

All data generated or analysed during this study are included in this published article and its supplementary information files. Custom R scripts are available upon request.

### Competing interests

The authors declare that they have no competing interests.

### Funding

This research was supported by the Innovation Technology Commission of Hong Kong SAR [grant number ITS113/15FP]; and Shenzhen Science and Technology Innovation Commission [grant number JCYJ20170818113656988].

### Authors’ contributions

H.X. conceived and initiated the study; X.L. collected the data and performed computational analysis; and H.X. and X.L. wrote the paper. Both authors read and approved the final manuscript.

## Acknowledgements

We thank Dr. Taobo Hu and Dr. Siu-Kin Ng for expert advice, and Dr. Zhenggang Wu for server management.

## Notes

### Competing Interest Statement

The authors have declared no competing interest.

