## Supplementary File S1. Supplementary figures S1-S12, captions of Supplementary Tables S1-S11, and description of Supplementary Dataset S1. for "Genetic-variant hotspots and hotspot clusters in the human genome facilitating adaptation while increasing instability"

**Contents:**

- Supplementary Figures S1 to S12
- Captions of Supplementary Tables S1 to S11
- Description of Supplementary Dataset S1
- Supplementary reference

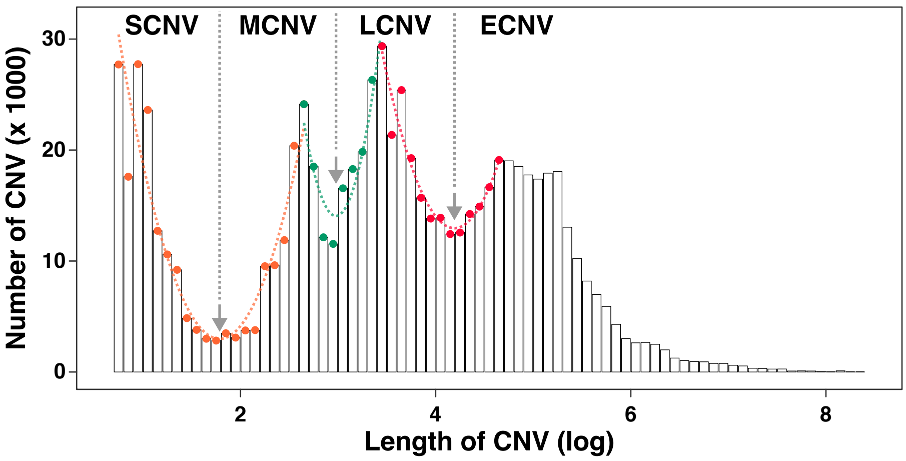


**Supplementary Figure S1. Size-classification of germline CNV regions retrieved from dbVar database.** Based on the $log_{10}$-transformed length distribution, CNVs are classfied into SCNV, MCNV, LCNV and ECNV based on the critical points (indicated by arrows) of the polynomial regression curves colored in orange, green and red respectively.

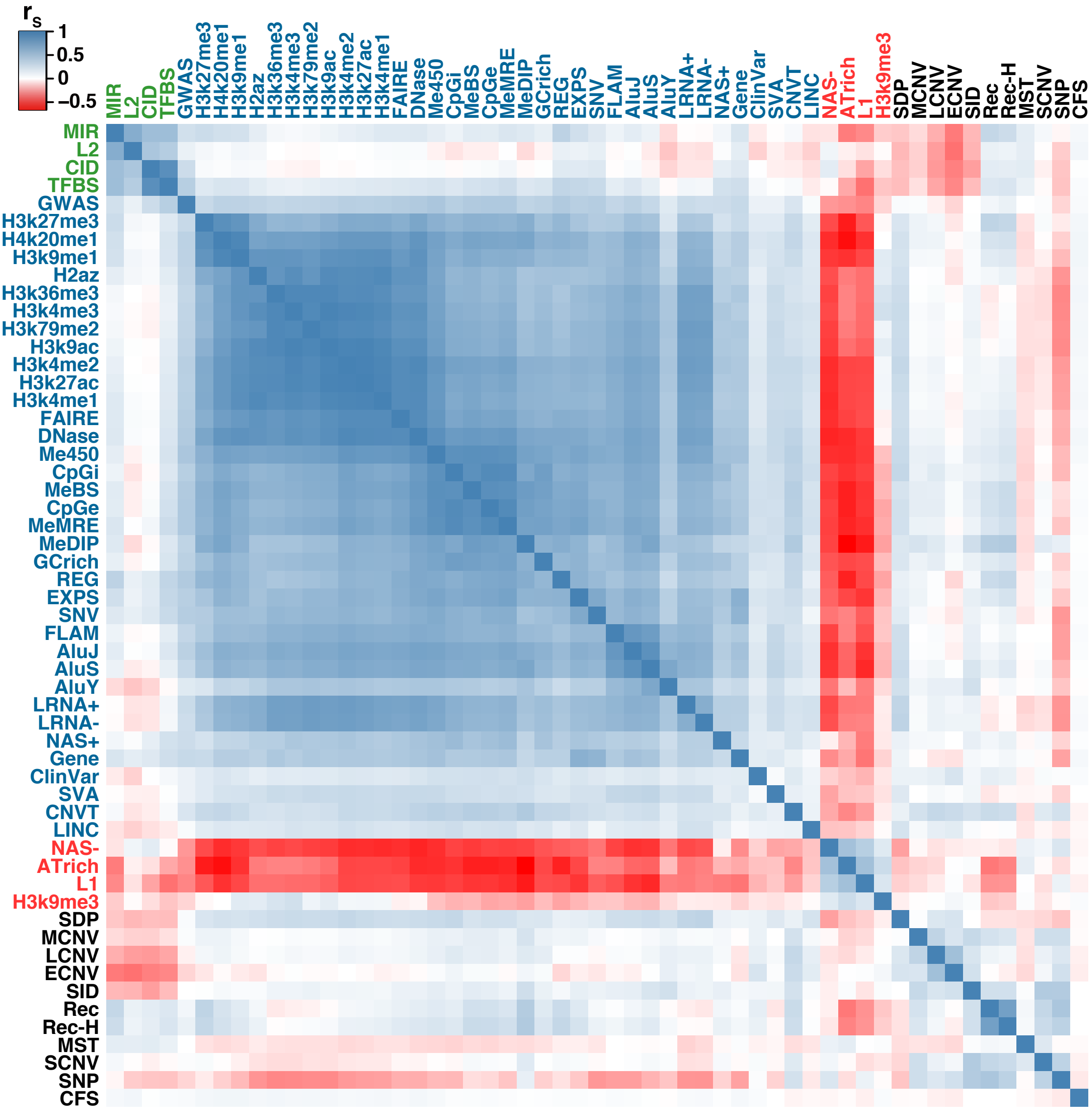


**Supplementary Figure S2. Heat map of pairwise Spearman's correlation coefficients ($r_S$) among different genomic features and markers in 500-kb non-overlapping autosomal windows.** The pairwise correlation coefficients are expressed in accordance to the red-blue thermal scale with blue square representing positive $r_S$ and red square representing negative $r_S$. The nature and clustering patterns of these genomic features enable their classification into the Genic zone (blue), Proximal zone (green), Distal zone (red) or marker (black) features. Markers including GWAS, CNVT and ClinVar which are allocated to the Genic zones on account of their propensities for positive co-localization with features in these zones.

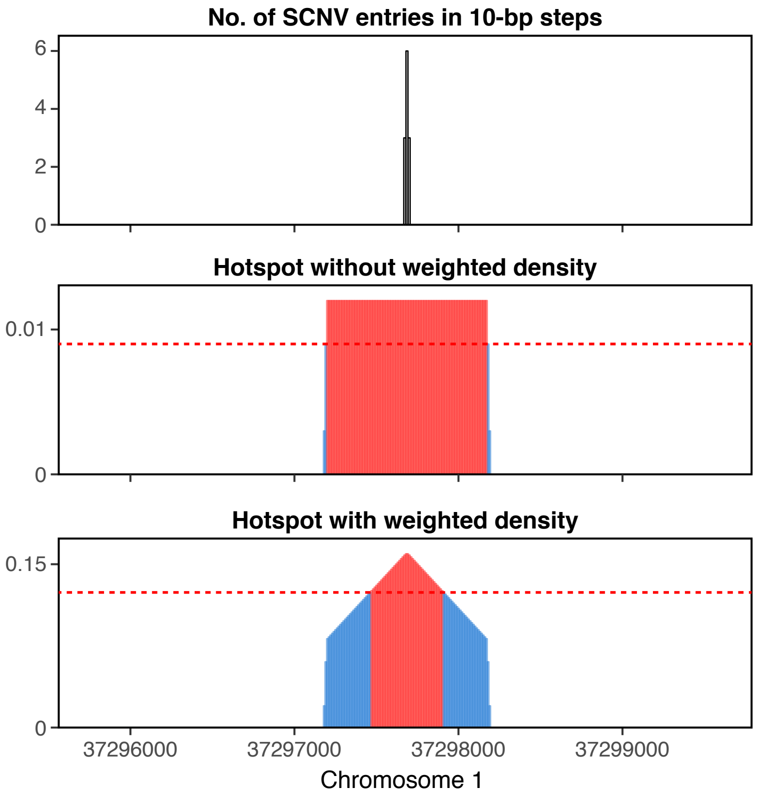


**Supplementary Figure S3. Refinement of hotpsot boundary by weighted density.** Top panel: local concentration of SCNV breakpoints in three 10-bp bins on chromosome 1. Middle panel: hotspot identified using SCNV density as criterion. Bottom panel: hotspot region identified using weighted SCNV density as criterion in sliding windows. In middle and bottom panels, windows with SCNV densities above threshold (red dashed line) are identified as hotspot (red bins) in sliding windows. With the weighting scheme described in Methods, sliding windows overlapping with the SCNV peaks are assessed with different levels of weighted SCNV density based on the location of SCNV peaks in each sliding window, enabling thereby the refinement of hotspot boundary.

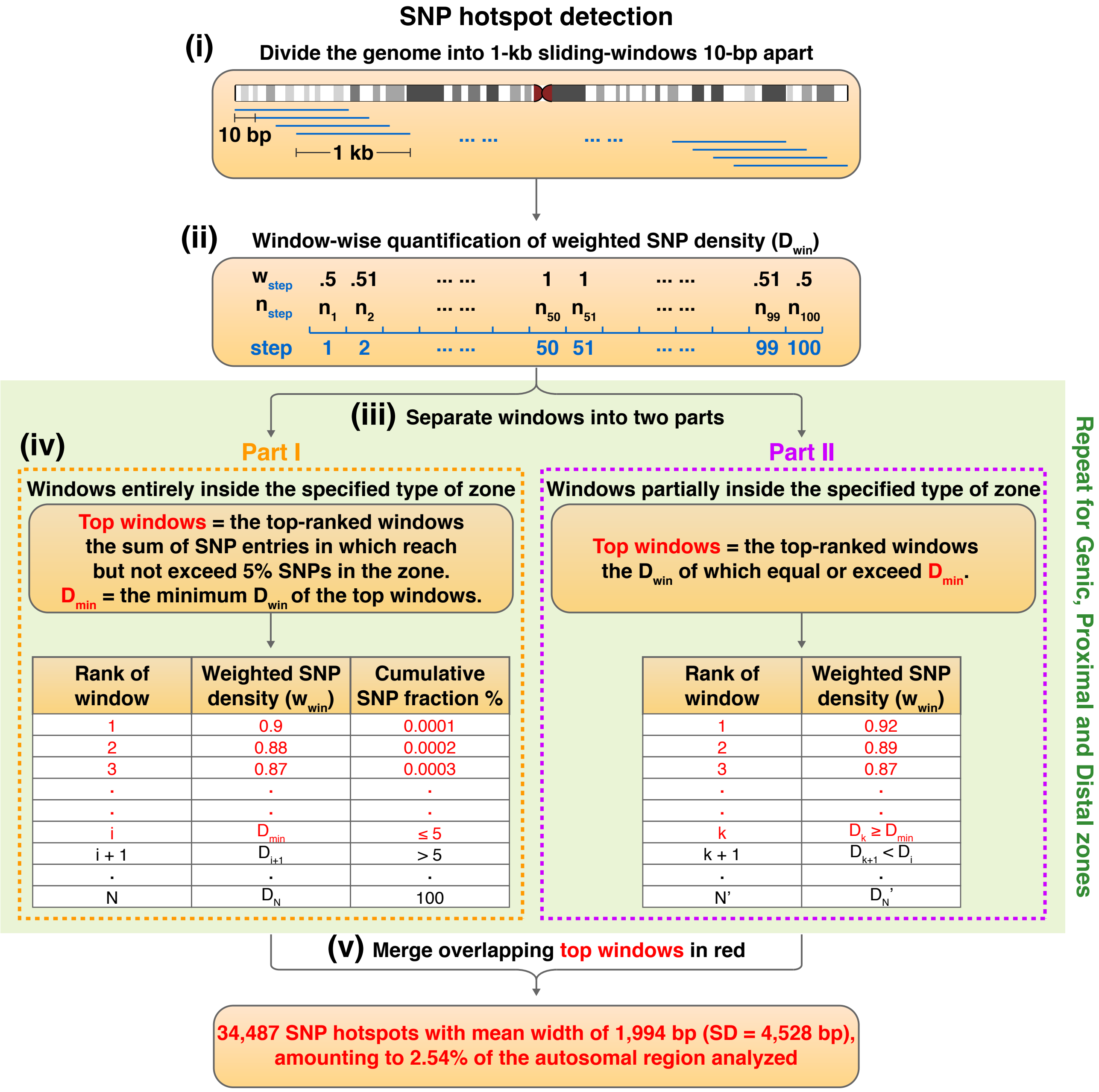


**Supplementary Figure S4. Flowchart of protocol for identifying SNP hotspots.** The protocol is described in "Density-based genetic-variant hotspots determined by weighted sliding windows" in Materials and Methods.

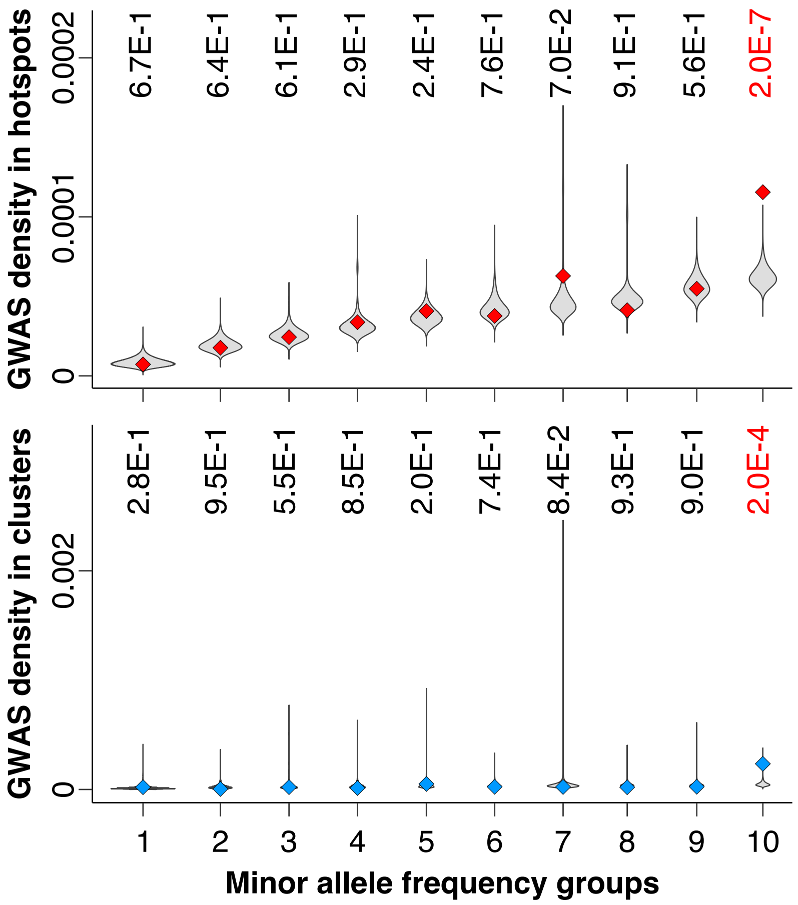


**Supplementary Figure S5. Density of GWAS-identified SNPs in ten groups of hotspots and clusters ranked based on minor allele frequency from low (group 1) to high (group 10).** Densities in simulated regions with matching minor allele frequencies are shown by the grey violin plots. A combined result of the ten groups is shown in Fig. 3d.

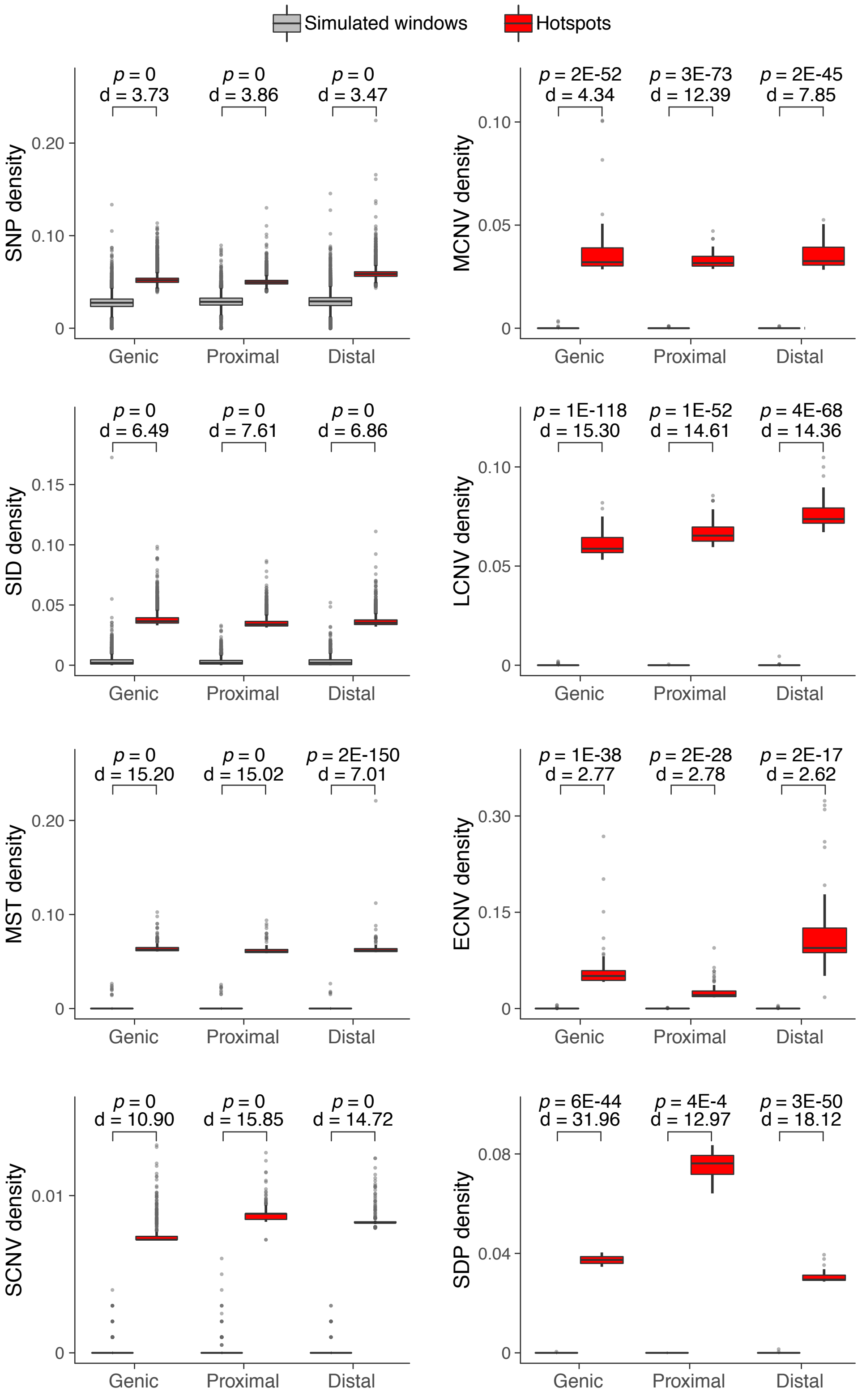


**Supplementary Figure S6. Comparison of GV densities in hotspots (red boxplots) and in simulated windows with matching number and size of the hotspots (grey boxplots).** The *p*-value and effect size (Cohen's d) are shown above each pair of boxplots.

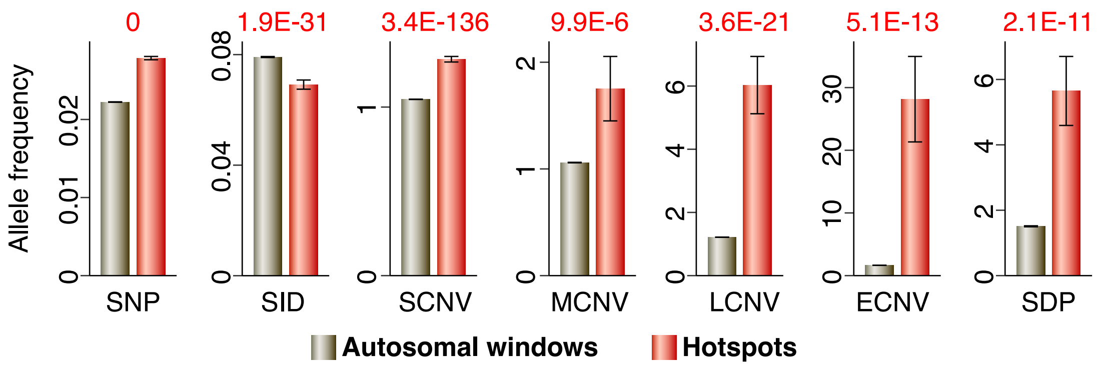


**Supplementary Figure S7. Comparison of GV allele frequencies in hotspots (red bars) and in total autosomal windows (grey bars) binned by the average size of GV hotspots (1,887 bp).** The *p*-value is shown above each pair of bars. The allele frequencies of SNP and SID is retrieved from 1000 Genomes Project. Recurrency is employed as allele frequency for SCNV, MCNV, LCNV, ECNV and SDP breakpoints. MST lacking allele frequency information is not shown in this figure.

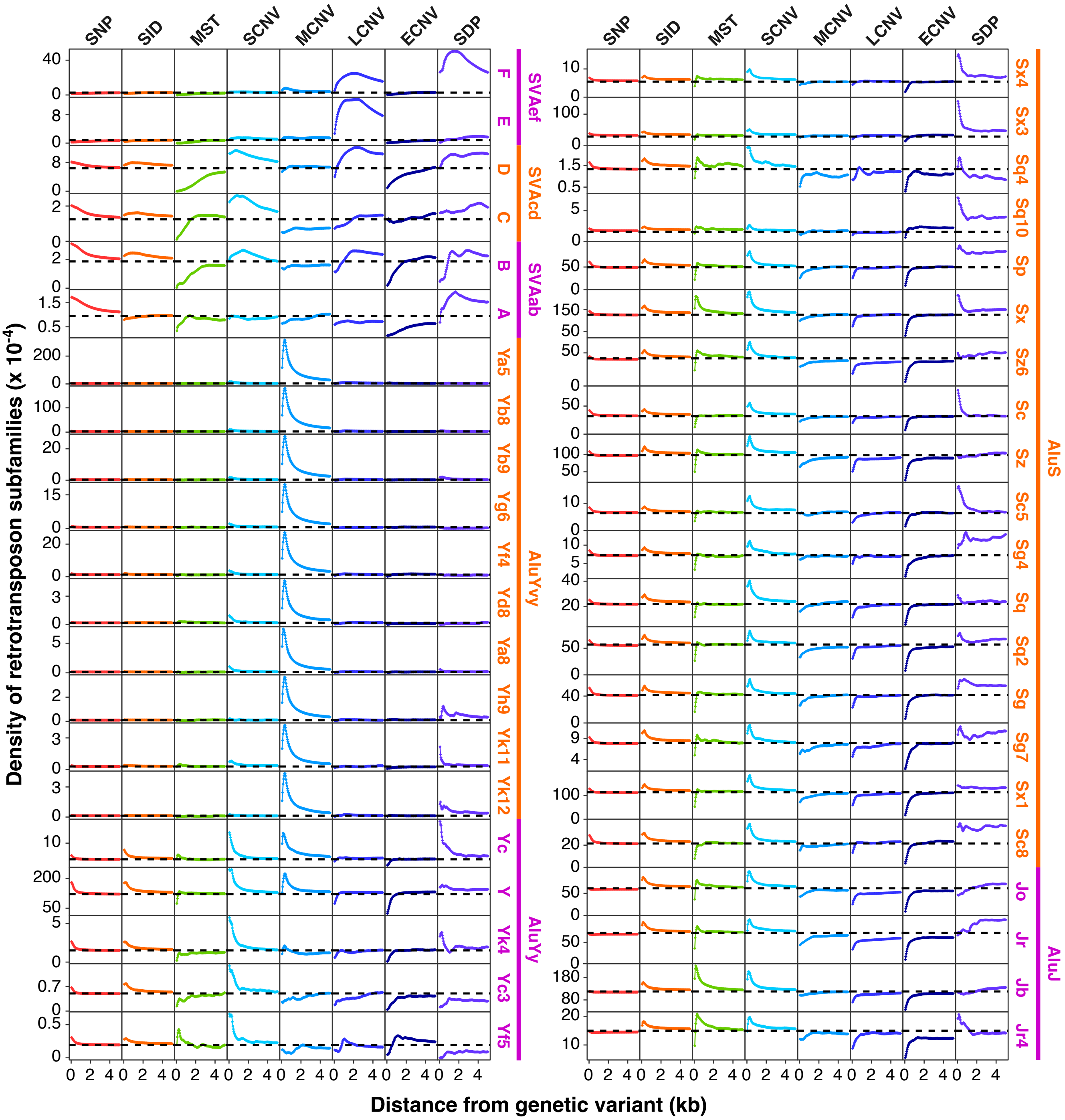

**Supplementary Figure S8. Density distribution of SVA and Alu subfamilies near eight kinds of genetic variants.** Cumulative density of 42 SVA and Alu subfamilies within ± 5-kb of SNP, SID, MST, SCNV, MCNV, LCNV, ECNV and SDP entries (in 50-bp increments). The horizontal dashed lines indicate autosomal averages of the retrotransposon subfamilies. The 6 SVA subfamilies and 15 AluY subfamilies shown in the left panel are merged into 5 larger groups in Fig. 4.

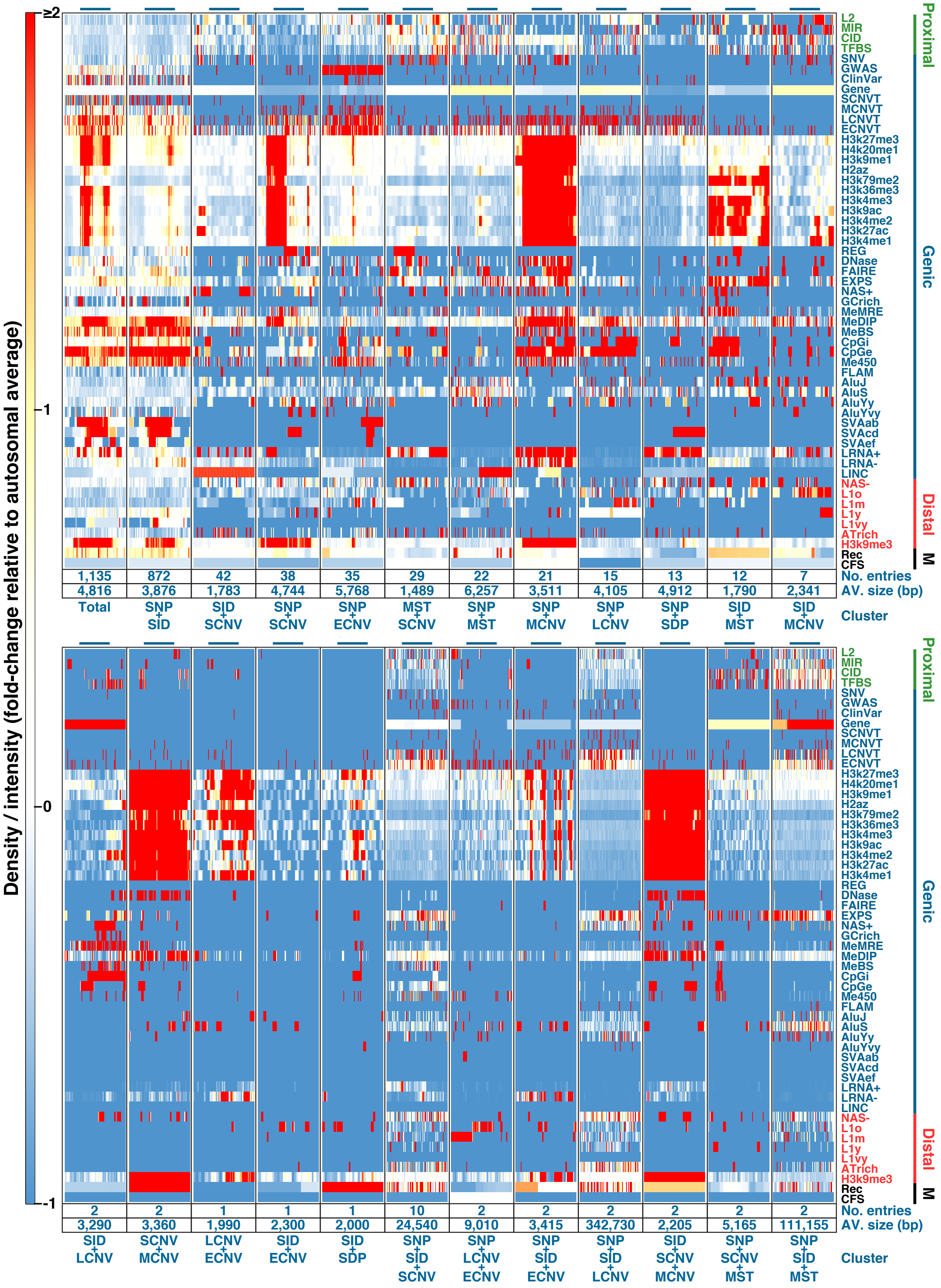


**Supplementary Figure S9. Genomic-feature contents of total (first column in upper panel) and 23 kinds of hotspot clusters with distinct hotspot-compositions.** The numbers and average sizes of each kind of cluster are indicated in the bottom two rows of each column. Fold-changes of the density/intensity of the genomic features are expressed similar to those in Fig. 8.

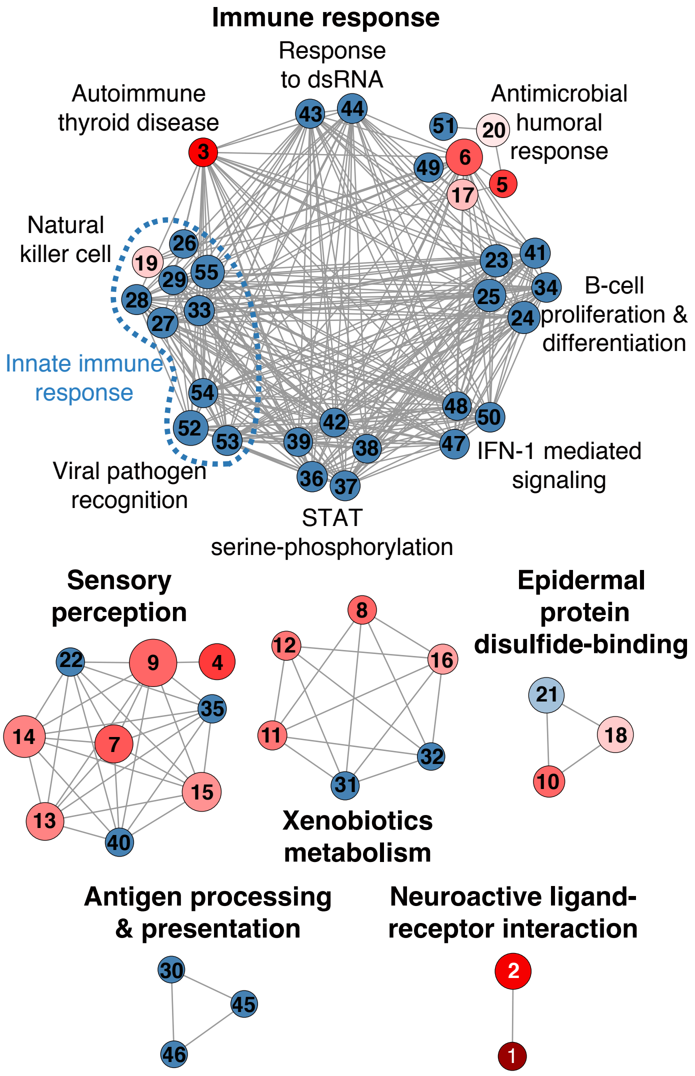


| ID | Pathway | Fold | ID | Pathway | Fold |
|---|---|---|---|---|---|
| 1 | KEGG:05033 | 15.72 | 29 | GO:0002323 | -1.00 |
| 2 | KEGG:04080 | 4.22 | 30 | GO:0002475 | -1.00 |
| 3 | KEGG:05320 | 3.56 | 31 | GO:0006067 | -1.00 |
| 4 | GO:0007608 | 2.19 | 32 | GO:0006069 | -1.00 |
| 5 | GO:0009620 | 2.14 | 33 | GO:0030101 | -1.00 |
| 6 | GO:0006959 | 1.83 | 34 | GO:0030183 | -1.00 |
| 7 | GO:0050906 | 1.74 | 35 | GO:0050912 | -1.00 |
| 8 | GO:0034308 | 1.69 | 36 | GO:0033135 | -1.00 |
| 9 | GO:0007606 | 1.67 | 37 | GO:0033138 | -1.00 |
| 10 | GO:0018149 | 1.64 | 38 | GO:0033139 | -1.00 |
| 11 | KEGG:00830 | 1.43 | 39 | GO:0033141 | -1.00 |
| 12 | KEGG:00982 | 1.39 | 40 | GO:0050913 | -1.00 |
| 13 | GO:0050907 | 1.30 | 41 | GO:0042100 | -1.00 |
| 14 | GO:0051606 | 1.30 | 42 | GO:0042501 | -1.00 |
| 15 | GO:0009593 | 1.13 | 43 | GO:0043330 | -1.00 |
| 16 | KEGG:05204 | 0.95 | 44 | GO:0043331 | -1.00 |
| 17 | GO:0019730 | 0.58 | 45 | GO:0048003 | -1.00 |
| 18 | GO:0009913 | 0.30 | 46 | GO:0048007 | -1.00 |
| 19 | GO:0001906 | 0.21 | 47 | GO:0060337 | -1.00 |
| 20 | GO:0032103 | 0.18 | 48 | GO:0060338 | -1.00 |
| 21 | GO:0030216 | -0.49 | 49 | GO:0061844 | -1.00 |
| 22 | GO:0001580 | -1.00 | 50 | GO:0071357 | -1.00 |
| 23 | GO:0046651 | -1.00 | 51 | GO:0071624 | -1.00 |
| 24 | GO:0032943 | -1.00 | 52 | KEGG:04621 | -1.00 |
| 25 | GO:0070661 | -1.00 | 53 | KEGG:04622 | -1.00 |
| 26 | GO:0002228 | -1.00 | 54 | KEGG:04623 | -1.00 |
| 27 | GO:0002285 | -1.00 | 55 | KEGG:04650 | -1.00 |
| 28 | GO:0002286 | -1.00 | | | |

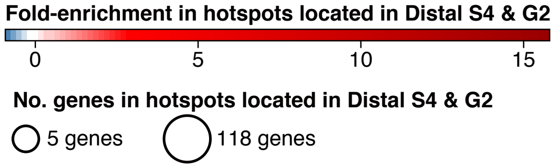


**Supplementary Figure S10. Enrichment Map for Distal-zone-enriched genes annotated using g:Profiler based on Gene Ontology biological process and KEGG.** Each circular node represents a pathway (with 3 to 350 genes) significantly enriched in Distal zones with Benjamini-Hochberg false discovery rate < 0.05, and the node size is proportional to the number of Distal-zone genes belonging to the pathway, as illustrated by the two circle signs of '5 genes' and '118 genes'. The pathway IDs inside the nodes correspond to those given in the table on the right. Pathways are connected by a grey edge when they share ≥ 50% genes. Enrichment of these pathway genes in the GV hotspots, represented by the red-blue thermal scale and shown in the 'Fold' column of the table, is estimated as fold-change of the fraction of the genes relative to the fraction of 1-kb windows that overlap with GV hotspots located in the S4- and G2-phase windows of Distal zones. Supplementary Table S11 gives the description of all the pathways shown in the figure.

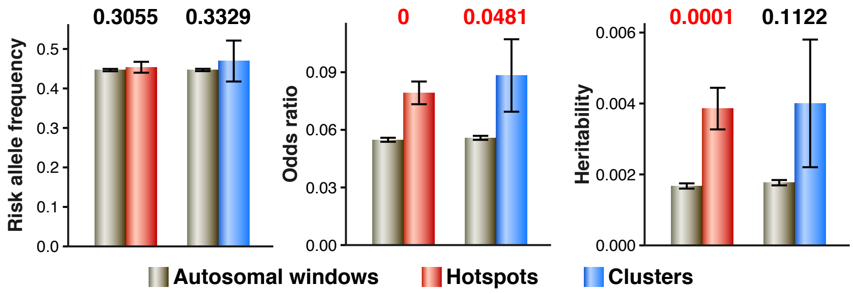


**Supplementary Figure S11. Comparison of risk allele frequency, odds ratio and heritability of GWAS-identified SNPs located within total autosomal windows (grey bars) relative to those located within hotspots (red bars) or clusters (blue bars).** The *p*-value are shown above each pair of bars. The risk allele frequency, odds ratio and heritability of GWAS Catalog SNPs (file date 2020.01.01) are estimated using GWEHS application (López-Cortegano and Caballero 2019) with all default settings and a prevalence value of 1%.

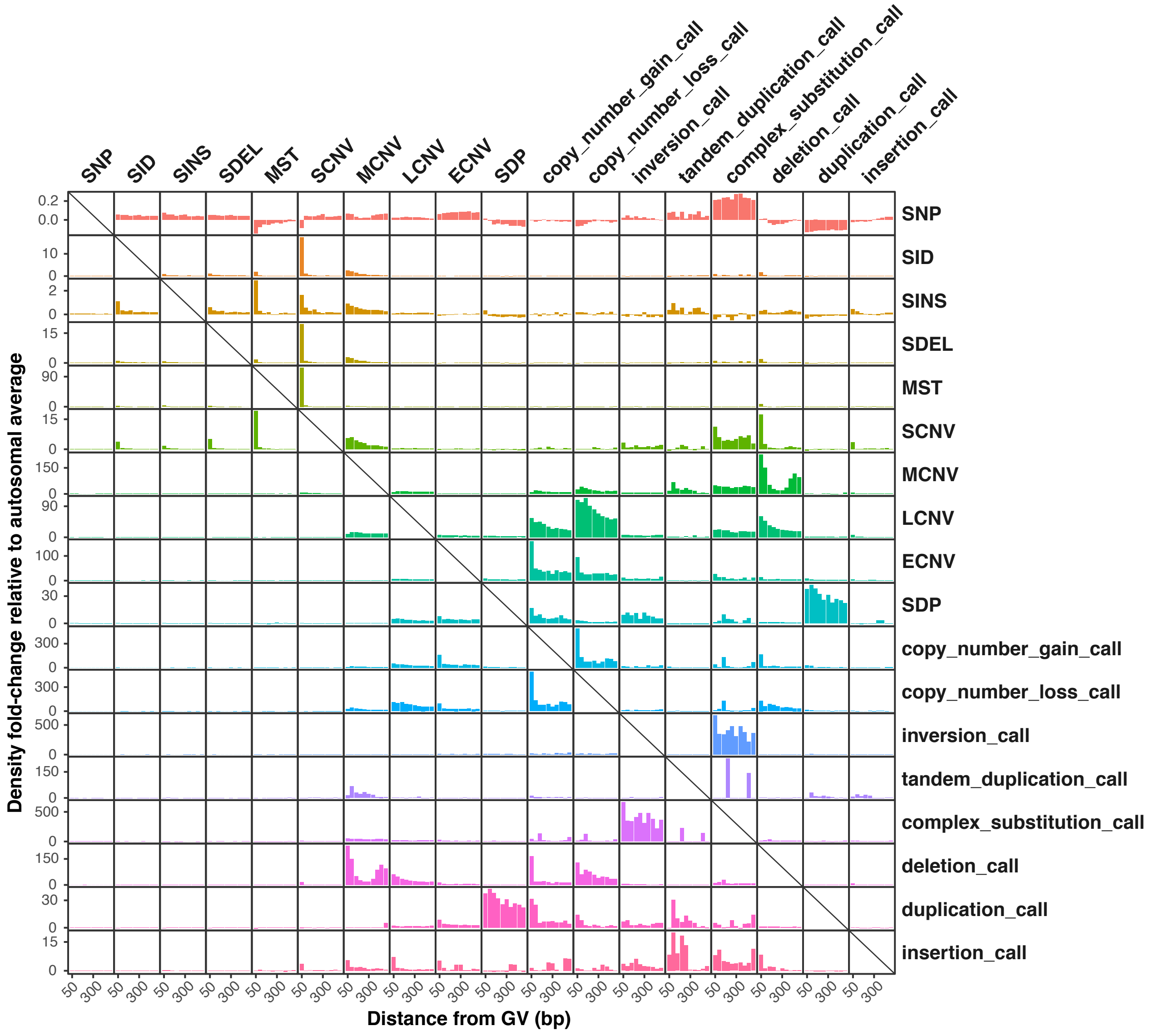


**Supplementary Figure S12. Co-localization among different kinds of genetic variants.** Enrichment of y-axis GVs at different distances from x-axis GVs, ranging from 0 bp to ± 500 bp in 50-bp increments. Enrichment of each y-axis GV is expressed by its density fold-change relative to its autosomal average density. Small indels (SID) are further separated into small insertions (SINS) and small deletions (SDEL). In addition to the eight kinds of GVs employed in hotspot detection, the variations analyzed in figure also include the 'calls' of other eight kinds of structrual variations given in dbVar database.

### Captions of Supplementary Tables

Supplementary Table S1. Information on genomic features and markers.

Supplementary Table S2. Properties of genetic-variant hotspots and hotspot clusters.

Supplementary Table S3. Properties of selection hotspots.

Supplementary Table S4. List of cluster-containing genes.

Supplementary Table S5. Estimated ages of retrotransposons.

Supplementary Table S6. Disease-related variants in GV hotspots and clusters.

Supplementary Table S7. Overlap of genetic-variant hotspots or clusters with recombination hotspots.

Supplementary Table S8. Autosomal sequences (in base pairs) in different genomic zones and replication phases that overlap with hotspots or clusters.

Supplementary Table S9. Enrichment and selection status of hotspots in seven immune system gene loci.

Supplementary Table S10. Genomic-feature distributions in genomic regions belonging to different replication-phase segments among 500-kb Genic, Proximal and Distal zones.

Supplementary Table S11. Functional annotation of Distal-zone genes.

### Descriptions of Supplementary Dataset

Supplementary Dataset S1. Information of genetic-variant hotspots, hotspot clusters, selection hotspots and replication-phase segments. Genomic coordinates are given in one-based format in human genome assembly hg19.
